# SpotDMix: informed mRNA transcript assignment using mixture models

**DOI:** 10.64898/2025.12.15.693918

**Authors:** Kasper Smeets, LuukW. Hesselink, Emmanuel Marquez-Legorreta, Greg M. Fleishman, Mark Eddison, PaulW. Tillberg, Misha B. Ahrens, Bernhard Englitz

## Abstract

Unveiling the genetic profiles of spatially distinguished cells is an important aspect in many areas of brain research, as the genetic identity contains information about a cell’s physiological properties and internal state. On top of this, knowledge of the genetic details of each cell can reveal structural organization within tissue. As image-based spatial transcriptomics moves toward applications in tissues with dense cellular packing, accurate assignment of detected mRNA transcripts (“spots”) to correct segmented cells becomes increasingly difficult, rendering simple methods insufficient with many incorrect assignments to neighboring cells. Here we introduce SpotDMix, a statistical model for assigning spots to cells by modeling spots as coming from a mixture model of distributions matching segmented cell shapes, with assignment probabilities and shape parameters optimized using the Expectation Maximization algorithm. Performance is assessed and compared against several simple methods in various scenarios on both surrogate data and larval zebrafish data. In all tested scenarios SpotDMix outperforms the simple methods on all evaluated metrics, including individual transcript assignment accuracy, total assigned number of spots per cell error and cell type classification. Further, SpotDMix produces a higher degree of exclusivity between genes which are known to not or rarely co-express.

## INTRODUCTION

The function of a cell is largely determined not by the DNA but by the expression of genes and production of proteins within the cell. The genetic expression of spatially distinguished cells not only gives information on cell properties and states but it also reveals important spatial structures within a tissue (Crosetto et al., 2015; Moffitt et al., 2022). Image-based spatial transcriptomics is an emerging field for imaging mRNA transcripts or other genetic markers while preserving their original spatial context (Moffitt et al., 2022). Fluorescent In-Situ Hybridization (FISH) methods such as osmFISH (Codeluppi et al., 2018), MERFISH (Chen et al., 2015b) or EASI-FISH (Wang et al., 2021) allow for detecting individual mRNA transcripts at subcellular resolution through fluorescent imaging, making it possible to map expression counts of selected genes to individual cells.

When combined with (ideally single-cell accurate) recordings of brain activity, for example from methods as multiMAP (Lovett-Barron et al., 2017; Marques et al., 2020) or light-sheet microscopy (Ahrens et al., 2013; Mahou et al., 2014), it becomes possible to study the complex link between a cell’s genetic profile and its functional or behavioral role (Bugeon et al., 2022; Lovett-Barron et al., 2017, 2020; Xu et al., 2020; Marquez-Legorreta et al., in prep.).

In FISH-based methods mRNA transcripts show up as bright diffraction-limited “spots” that can be localized using image processing methods. Detected spots for different genes can then be assigned to segmented cells in order to obtain transcript counts and estimates of the genetic profiles of individual cells. Existing FISH-based methods have mostly been applied to mouse brain areas with cell segmentation performed based on cytoplasmic markers such as poly(A) (Codeluppi et al., 2018; Chen et al., 2015b; Wang et al., 2018) or cytoDAPI (Wang et al., 2021, Wang et al., 2018). Cytoplasm-based segmentation is ideal for spot assignment when cells are spatially separated. While mRNA transcript production occurs in the nucleus of the cell, transcripts are transported to the cytoplasm where translation and protein creation occurs, resulting in the majority of mRNA transcripts being localized inside the cytoplasm and outside of the nucleus.

Unlike in mice, cells in the brain of larval zebrafish are densely packed with nuclei taking up the vast majority of the volume of a cell (Fig. 1C), leaving no detectable boundaries between cells when using cytoplasmic markers and thus making cytoplasm-based segmentation practically impossible. Instead, cell segmentation can be done using a nuclear marker such as DAPI at the cost of losing information about the cytoplasm. As most of the spots are localized outside of the nuclei, it becomes challenging to correctly assign the spots to the segmented cells due to the high packing density and the lack of cell boundary markers (Fig. 1G). As will be shown, using simple methods such as assigning spots to the nearest nucleus often fails as spots are falsely assigned to neighboring cells instead.

**Figure 1:**
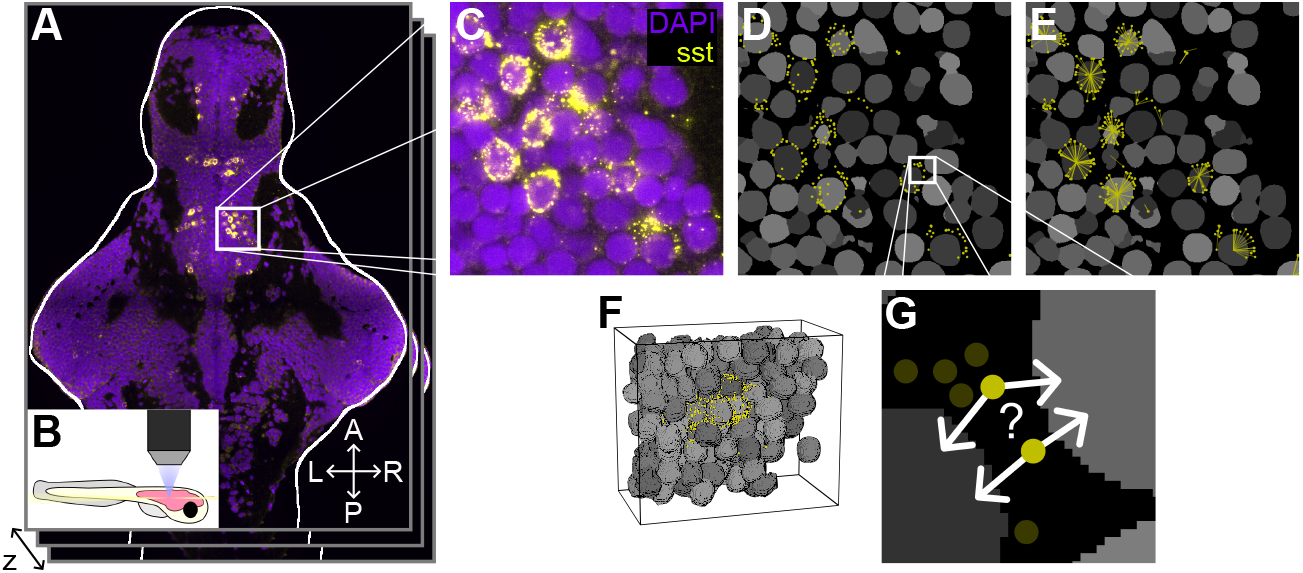
**A)** Combined raw fluorescent larval zebrafish WARP data of DAPI (purple, staining nuclei) and sst (yellow, somatostatin mRNA transcripts), highlighting a single slice. L/R: Left/Right. A/P: Anterior/Posterior. **B)** Depiction of the setup used for measuring the WARP data. **C)** Enlarged view of the box in A, highlighting a section of the thalamus. Diffraction limited spots are often localized in between the dense network of cells. **D)** Processed data corresponding to the box in B showing segmentation (gray) with detected spots (yellow). **E)** *Hard* assignments from SpotDMix follow the visually clear radial patterns of spots. **F)** 3D render of a small section in the thalamus showing the complex and dense structure of segmentation and sst spots. **G)** Enlarged view of the box in D. As spots are typically localized in between segments, from the perspective of a single spot it is often impossible to determine to which cell it should be assigned.

One method for assigning spots which lie outside of the segmentation is Sparcle (Prabhakaran, 2022), which works by estimating the genetic profile of each cell and has been shown to work well on mouse data. This method however requires the measurement of mRNA transcripts from many genes, in order to get accurate genetic clusters, as well as for enough spots to be localized within the segmentation to get good initial cell type clusters. Even then, spots are assigned to the closest cell matching their cell type.

Several other methods exist that do not require prior segmentation of cells such as Baysor (Petukhov et al., 2022) and SSAM (Park et al., 2021), among many others (Andersson et al., 2024; Partel and Wählby, 2021; Mages et al., 2023), which can directly cluster spots from many genes into cell types and in the case of Baysor even generate segmentation from the gene expression data itself. However, these methods either rely on high mRNA density within cells in order to estimate cell locations or on having enough genes to accurately identify cell types within the data. Several of the methods mentioned here are also only applicable to identify cell types and spatial structures as they do not produce genetic expression counts per cell, making them unsuitable for direct use in multi-modal analysis.

Here we introduce SpotDMix, a method for spot assignment that makes use of the visually observed spatial distributions of spots around cells seen in dense tissue. Spots are processed separately for each gene by modeling these spots as being drawn from a weighted mixture of radial spatial distributions around cells. Inspired by Gaussian Mixture Models (GMMs) (Reynolds, 2009), it uses the Expectation Maximization (EM) algorithm (Dempster et al., 1977; Bishop, 2006) to iteratively estimate the shape of the distribution, the relative weighting of each cell and the probability of each spot to belong to any specific cell. SpotDMix provides a robust and efficient statistical method for spot assignment to segmentation in dense tissue without the need for prior cell type identification, using expected spatial distributions of spots around cells. The performance of this method is evaluated on a data set from larval zebrafish (Marquez-Legorreta et al., in prep.), both on surrogate data generated from the available cell segmentation as well on the actual detected spots, to assess and compare its accuracy and classification performance to non-informed methods.

## RESULTS

### A mixture model of spot locations

In the raw fluorescent data associated with a gene, where mRNA transcripts show up as diffraction limited spots, there is often a clear visual distinction between cells which express a certain gene and those which do not, where transcripts form a shell-like structure around the nucleus (Figure 1A-B). In essence, SpotDMix assigns spots by demixing spot locations as if drawn from defined spatial distributions around cells. A spatial probability distribution 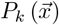 is defined around every cell *k*, specifying the expected locations of spots in the case that the cell would be positively expressing for a gene. The measured positions of all spots for a single gene is then seen as an observation from a weighted mixture of the individual cells:

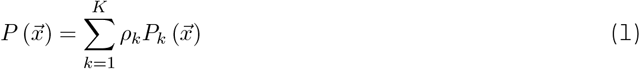

where *ρ*_*k*_ is the mixture weight corresponding to cell *k*, and *K* is the total number of cells (i.e. components).

The computational steps that make up SpotDMix are outlined in Figure 2A and are detailed below and in the Methods section. For the present problem setting, the chosen probability distributions are transformed radial distributions based on ellipsoidal fits of cells. Specifically, this is done by defining a radial gamma distribution in the space defined by the Mahalanobis distance around each cell:

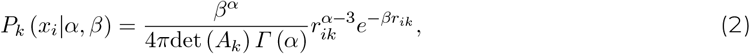

where

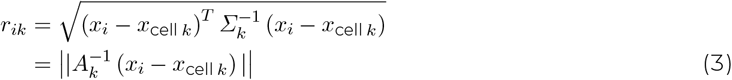

is the transformed (Mahalanobis) distance between the location of spot *i* and the center of cell *k*, and *A*_*k*_ is a transformation matrix defining the fitted ellipsoid.

**Figure 2:**
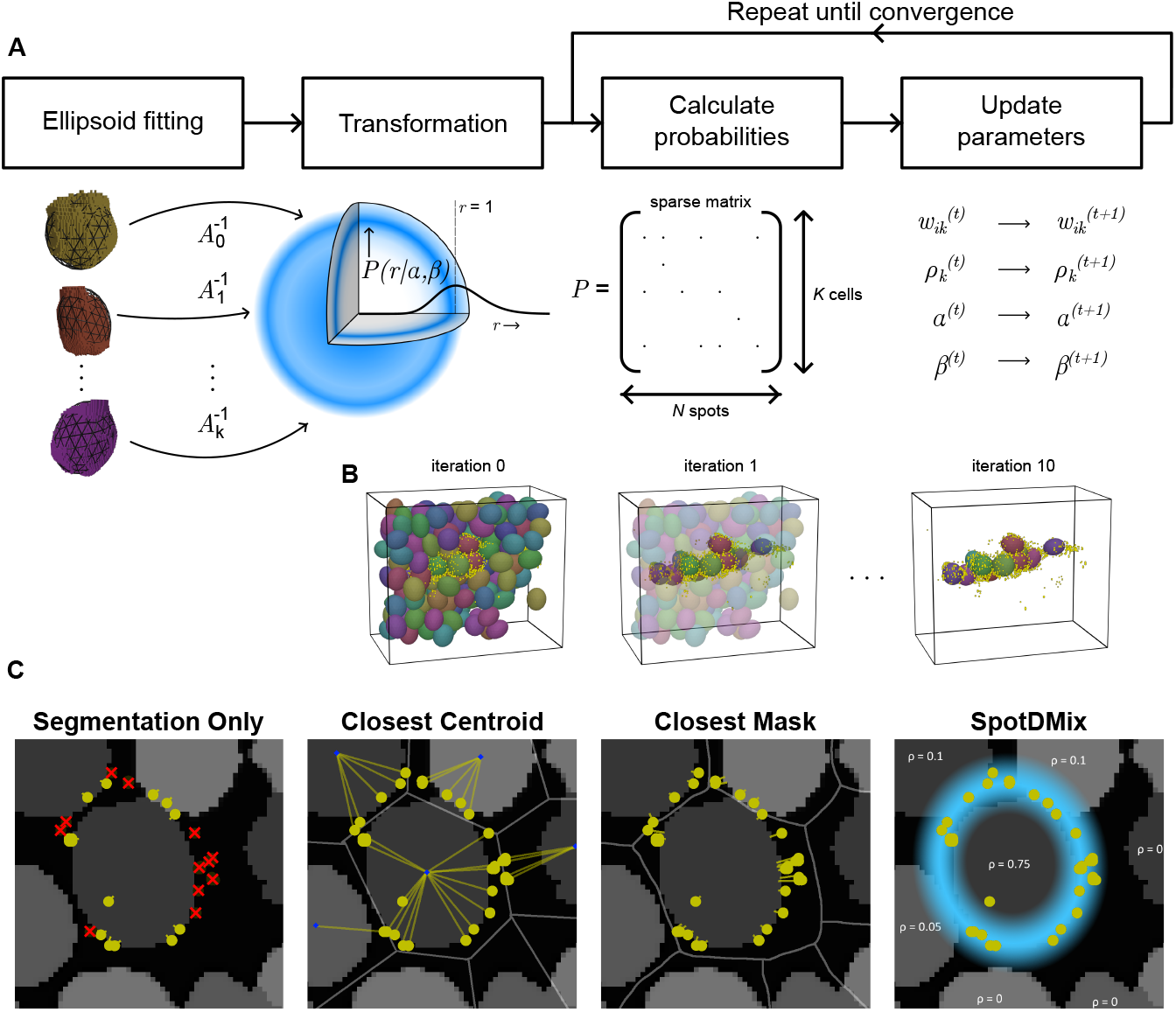
**A)** An overview of the steps involved in running SpotDMix. Ellipsoids are fit around segmented cells, and used for the transformation of spots to the radially symmetric space in which the probability density function is defined. In this space, a distance of *r* = 1 indicates that a spot is localized exactly on the border of the corresponding ellipsoid, with values less and greater than 1 meaning spots inside and outside the ellipsoid respectively. Probabilities are stored in a sparse matrix based on nearby cells for each spot. These probabilities are used to update the assignment probabilities, the relative weights and the shape parameters used for determining the probabilities. This process is repeated until the likelihood of the data is converged corresponding to a local maximum. **B)** Example iterations of SpotDMix corresponding to the area in Figure 1F, showing fitted ellipsoids and sst spots. The relative weight of each cell is indicated by the transparency of each ellipsoid and convergence is quickly reached with only cells with many spots around them retaining a high weight. **C)** The different methods evaluated and compared, showing example assignments for spots around a segment. Red crosses denote unassigned spots. Decision boundaries are indicated for Closest Centroid and Closest Mask in gray. *ρ* indicates relative weights of cells.

The shape parameters *α* and *β* determine the specific shape of the radial distribution and are iteratively updated during the assignment process. Crucially, the shape parameters are shared across all cells to ensure that cells without surrounding spots cannot extend their influence and ‘take’ spots from far away. Having shared parameters also allows cells to learn a common spatial distribution, as on a single cell level the number of spots can vary significantly. On top of this, the spot locations may not always perfectly adhere to expected distributions due to external factors such as natural variability, noise or improper segmentation.

The mixture weights *ρ*_*k*_ and the shape parameters *α, β* are iteratively found through the use of the EM algorithm. Given these parameters, the probability of spot *i* belonging to cell *k* (termed “*soft* assignments”) follows directly from applying Bayes’ rule:

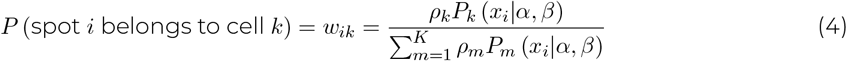

Initial assignment uncertainties between neighboring cells (as in Fig. 1G) are solved through the mixing weights *ρ*_*k*_, as the weights directly correlate to the number of spots that match the distribution *P*_*k*_ (*x*) around the cell. This breaks the symmetry and allows spots to be more likely to be assigned to a cell with more spots surrounding it.

SpotDMix is designed to quickly and efficiently run on large data sets without an explicit need for compute clusters (Suppl. Table 1), and convergence is quickly reached within only a few iterations (Fig. 2B). The probabilistic nature of this method allows for a multitude of ways to define the final spot assignments, two of which are explored here and are termed *soft* and *hard* assignments respectively. With *soft* assignments, spot counts per cell are exactly the sum of soft assignment probabilities, yielding a fair split in uncertain cases at the cost of not having quantized counts. For *hard* assignments, every spot is assigned to the cell to which it has the highest probability of assignment. Though this method seems a natural choice, as will be seen it may suffer slightly from overfitting when spots are equally well assigned to multiple cells.

### Model performance on surrogate data

As annotated genetic data is limited in zebrafish, surrogate data resembling physiological spots was generated based on a 3D volume of segmented cells ( ∼ 85 *×* 93 *×* 93 *µm*; 5040 cells; hindbrain). Two surrogate scenarios are considered to easily test performance for a variety of different factors, such as the sparsity of positive cells. Spot locations sampled from blurred dilated segmentation mask boundaries (see Methods).

The performance of SpotDMix is assessed and compared to three simple, non-informed, methods in the following two scenarios: i) Surrogate spots around sparsely selected cells, to compare individual spot assignment accuracy and positive/negative classification in an ideal case. ii) Surrogate spots around two neighboring but mutually exclusive populations, to evaluate how neighboring cell types can influence classification.

The methods used for the comparison are visually demonstrated in Figure 2C, and consist of: **Segmentation Only (SO)**: Assign only spots that overlap with segmentation, leave out the rest. **Closest Centroid (CC)**: Assign spots to the segment with the nearest centroid (center of mass). **Closest Mask (CM)**: Assign spots to the nearest (boundary of) a segment.

Closest Centroid is computationally the most efficient algorithm but is expected to lack in accuracy. It partially serves to find a lower bound on the performance of assignment methods. On the other hand, Closest Mask is computationally expensive to implement yet a natural choice for incorporating more information than is obtained from only using the spots overlapping with the segmentation. A more detailed description of the different methods is provided in the Methods section.

#### Sparse positive cells

Neuropeptides serve crucial roles within the nervous system, encompassing the regulation of hormone release, cell development, and specific behavioral functions (Herzog, 2012; Swanson and Sawchenko, 1983; Sandoval-Talamantes et al., 2020). Cells positively expressing neuropeptide-associated genes such as somatostatin (sst), neuropeptide Y (npy), oxytocin (oxt), and numerous others, are often only sparsely found throughout the brain (Lovett-Barron et al., 2020; Marques et al., 2020; Xu et al., 2020). While they can sometimes occur in small groups together, the vast majority of cells expressing neuropeptide-associated genes occur with considerable distances between positive cells.

In this simplest and first surrogate scenario, cells in the volume are randomly selected to be either positive or negative according to *P* (positive) = 0.03, with spots being generated around positive cells (Fig. 3A; Methods). All parameters were chosen to best match the distributions and statistics observed in the experimental data, such as the sparsity of and the number of spots around positive cells (see Methods).

**Figure 3:**
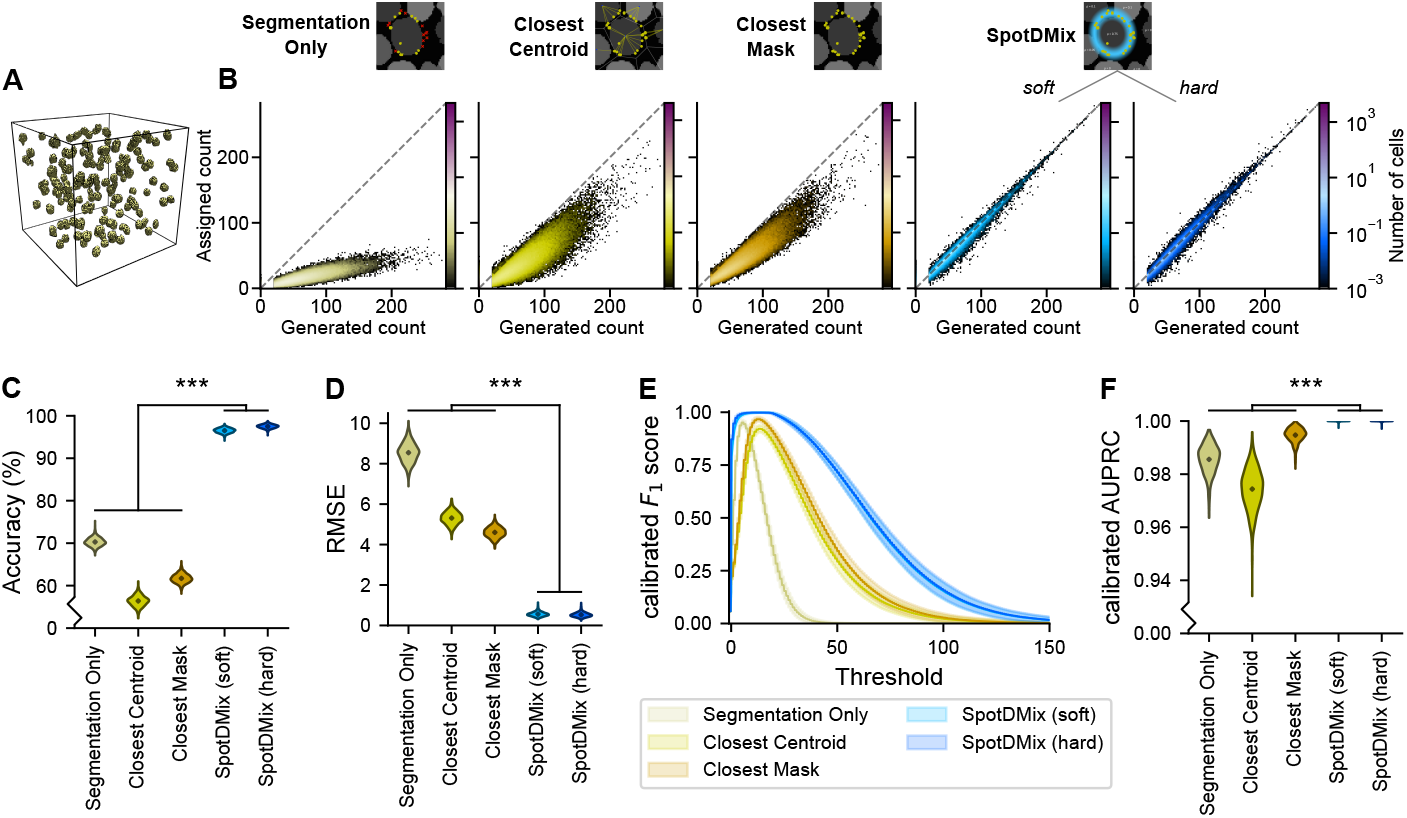
Performance of the different methods in surrogate scenario i), on **A)** sparsely selected cells in a excerpt of the hindbrain for spot generation. **B)** Heatmap comparing the number of spots generated around a cell to the number of spots that got assigned to each cell, for the different compared methods. Values indicate mean occurrence across 1000 repeated runs. Dashed lines indicates perfect assignments. The quality of the assignments is measured through the **C)** individual spot assignment Accuracy and the **D)** Root Mean Square Error between the generated and assigned number of spots per cell. The RMSE directly quantifies the deviation from the ideal line in B. **E)** calibrated *F*_1_ scores (mean *±* std), indicating the quality of classification as a function of the counts threshold. Low thresholds lead to over-classification and high thresholds lead to under-classification. **F)** The calibrated Area Under the Precision-Recall Curve (AUPRC) summarizes the classification performance, with a value of 1 indicating a perfect classifier. Violin plots (**C, D, E**) show the extent and distribution of the obtained values, with a diamond showing the mean value.

Here, both SpotDMix variants achieve a significantly higher individual transcript assignment accuracy than simple methods across 1000 repeated runs (all *p* ≤ 10^−307^, all effect sizes ≥ 33.0, pairwise Wilcoxon rank-sum tests, Suppl. Fig. 3) (Acc._*soft*_ = 96.5 *±* 0.6, Acc._*hard*_ = 97.5 *±* 0.5% versus Acc._SO_ = 70.3 *±* 1.0%, Acc._CC_ = 56.4 *±* 1.3%, Acc._CM_ = 61.7 *±* 1.1%; mean *±* standard deviation) (Fig. 3C). For most models the accuracy is defined as the percentage of spots that is correctly assigned to its corresponding ground truth cell. For the *soft* variant of SpotDMix, which does not have quantized assignment, the accuracy is defined as the mean probability of a correct assignment (see Methods).

Furthermore, gene counts obtained through SpotDMix are closer to the generated gene counts per cell, as measured through the Root Mean Square Error (RMSE). Both assignment variants of SpotDMix achieve a similar error (RMSE_*soft*_ = 0.56 *±* 0.11, RMSE_*hard*_ = 0.53 *±* 0.12), and both reach a significantly smaller error compared to all simple methods again for all runs (all *p* ≤ 10^−307^, all effect sizes ≥ 20.2, pairwise Wilcoxon rank-sum tests, Suppl. Fig. 3) (RMSE_SO_ = 8.5 *±* 0.5, RMSE_CC_ = 5.3 *±* 0.4, RMSE_CM_ = 4.60 *±* 0.26) (Fig. 3D). This distinction can also be visually observed through the heatmap comparing the number of generated spots to the number of assigned spots per cell (Fig. 3B), as the RMSE score directly summarizes the deviation from the ground truth diagonal line.

The Segmentation Only method stands out for having a large error in the assigned number of counts, which is largely due to how only 34.7 *±* 0.9% of spots in this scenario were assigned. However, despite the efforts to match the parameters for generating the spots as close as possible to what is observed in the experimental data, care should be taken when interpreting these results as the performance of the mask-based assignment methods strongly depends on the parameters used for generating this data (see Discussion).

Another crucial metric is that of classification performance. Independent of single-spot accuracy, how well can positive and negative cells be discerned by setting a threshold to separate positive from negative cells? The performance as a function of the chosen threshold is measured using the trade-off between precision (positive predictive value) and recall (true positive rate). A low threshold will have a high recall but low precision, and a high threshold will have a low recall and high precision. This trade-off is measured with the *F*_1_ score, which is the harmonic mean between precision and recall, and which exists for every chosen threshold (see Methods). It provides a visually clear comparison between methods and is useful in determining the optimal threshold to choose for classification. Here, a calibrated variant of precision (Siblini et al., 2020) is used to account for the variability in the number of positive cells in each run. This calibrated precision is also used for calculating the *F*_1_ score, obtaining the calibrated *F*_1_ (*cF*_1_) score.

From the *cF*_1_ curves for the different methods it is seen that both SpotDMix variants obtain a larger *cF*_1_ value for all thresholds compared to the simple methods (Fig. 3E), indicating a better classification performance regardless of the threshold chosen. In this scenario there is no discernable difference between the *cF*_1_ scores of the *soft* and *hard* SpotDMix variants.

*F*_1_ curves can look widely different for different methods, with peaks at different locations and different total widths depending on the distribution of assigned spot counts (e.g. Segmentation Only has a very narrow curve as less spots are assigned to cells). Both to remove the ambiguity for different optimal thresholds for different methods as well as to summarize the classification into a single scalar value, the calibrated Area Under Precision-Recall-Curve (cAUPRC) score is used (see Methods). This metric is similar to that of the more well-known metric of taking the Area Under the Curve (AUC) applied to ROC curves, with higher values indicating a better performance, but is better suited for cases with a large class imbalance as True Negative classifications do not count towards either the precision or the recall (Saito and Rehmsmeier, 2015; Flach and Kull, 2015; Siblini et al., 2020).

The cAUPRC scores for the different methods are summarized in Figure 3F, with both SpotDMix variants performing comparable (cAUPRC_*soft*_ = 0.999958 *±* 0.00022, cAUPRC_*hard*_ = 0.999962 *±* 0.00022) and significantly improving cell type classification compared to simple methods (all *p* ≤ 10^−307^, all effect sizes ≥2.7, pairwise Wilcoxon rank-sum tests, Suppl. Fig. 3) (cAUPRC_*SO*_ = 0.986 *±* 0.006, cAUPRC_*CC*_ = 0.974 *±* 0.008, cAUPRC_*CM*_ = 0.9947 *±* 0.0027). Despite the higher accuracy for the assigned spots from Segmentation Only, the more computationally intensive method of Closest Mask achieves a higher average classification performance in this scenario.

#### Large populations of neighboring cell types

Whereas neuropetides typically express sparsely, other genes express in distinct larger spatial populations. Examples include vglut2a for excitatory cells, gad1b for inhibitory cells (Appelbaum et al., 2009) (Fig. 5D) and pvalb7 for cells in the cerebellum (Yanicostas et al., 2012). These spatial structures of particular cell types often border with other populations of cell types without overlap between the two. This mutual exclusivity is evident in pairs such as gad1b and vglut2a, representing inhibitory and excitatory cells respectively, or gad1b and cx43, which distinguish (inhibitory) neurons from glial cells (Fig. 5D).

Cell type classification is most challenging at the boundaries between these regions of mutually exclusive cell types, as transcripts of both genes can occur close together. To further complicate matters, dense populations of single cell types are a challenging area for accurate spot assignment. As a large portion of the produced mRNA transcripts lie exactly in between the two or more positively expressing cells, it becomes impossible to confidently assign transcripts even for humans.

Scenario ii) consists of two continuously-stretching populations of cell types, with locations of positively labeled cells generated using Perlin noise patterns (see Methods, Fig. 4B). Spots are generated and processed independently for the two cell types, obtaining two separate spot counts per cell, and making it possible to assess to what extent the performance of the different methods is affected by having neighboring positive cells, and to what extent the different methods are able to retrieve this split in cell types.

**Figure 4:**
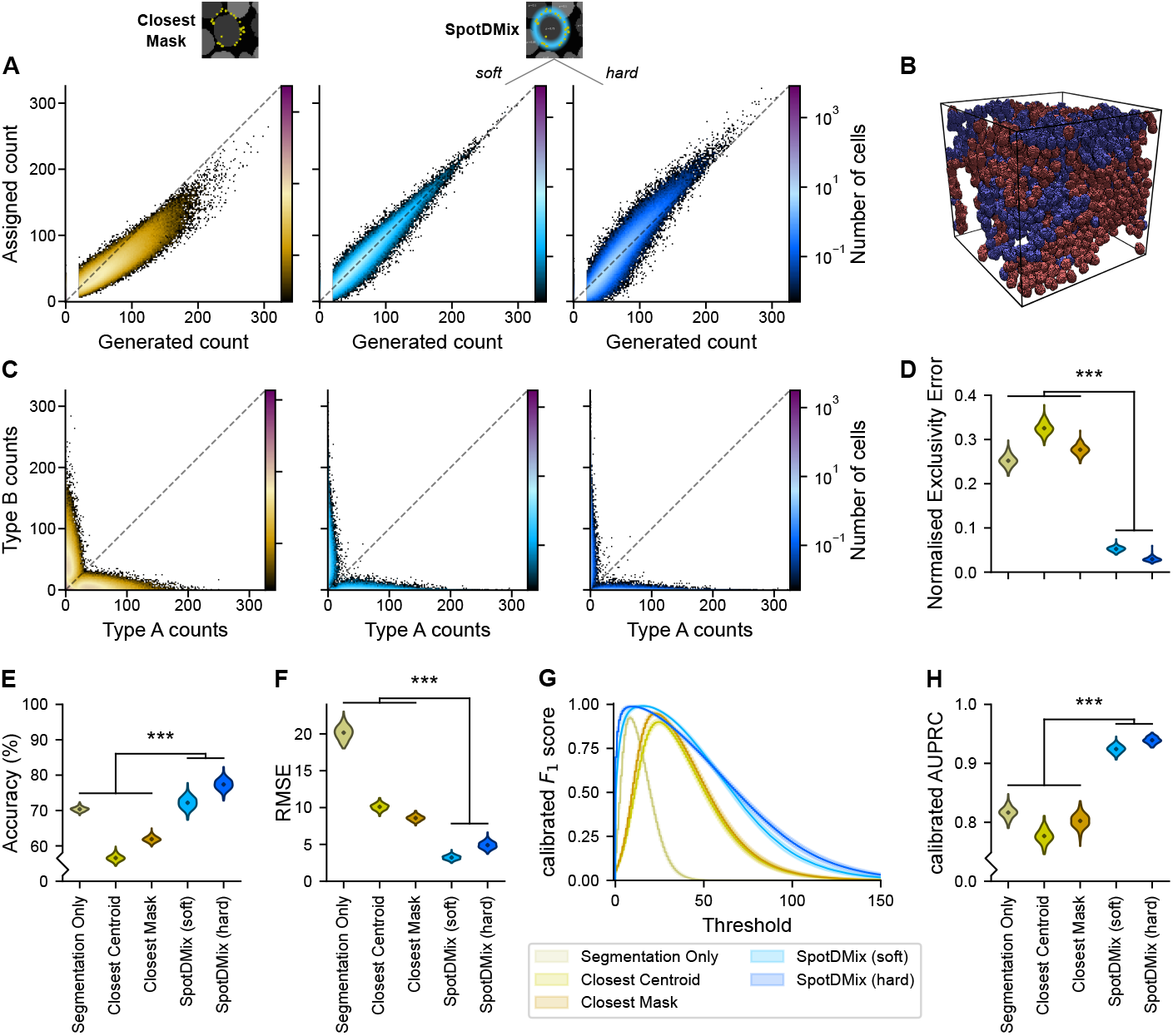
Performance of the different methods in surrogate scenario ii), consisting of two closely neighboring mutually exclusive populations of cell types. **A)** Same as in Fig. 3B, but only highlighting Closest Mask and SpotDMix. Results of the 200 runs are pooled for both cell types as the distributions they follow are symmetric. **B)** 3D render showing cells selected as positive for type A (red) and type B (blue) respectively, based on a Perlin noise pattern. **C)** Assigned counts for spots from type A and type B. As the two cell types were generated as being mutually exclusive, an ideal assignment scenario would correspond to no values beyond the axes themselves. **D)** The deviation from this ideal scenario is measured using the normalized Exclusivity Error (Methods), which relates to the spread observed in the plots in C. **E, F, H)** Same as in Figure 3, with results combined from both cell types. **G)** Mean calibrated *F*_1_ curve between one-v-all classification for both cell types separately.

**Figure 5:**
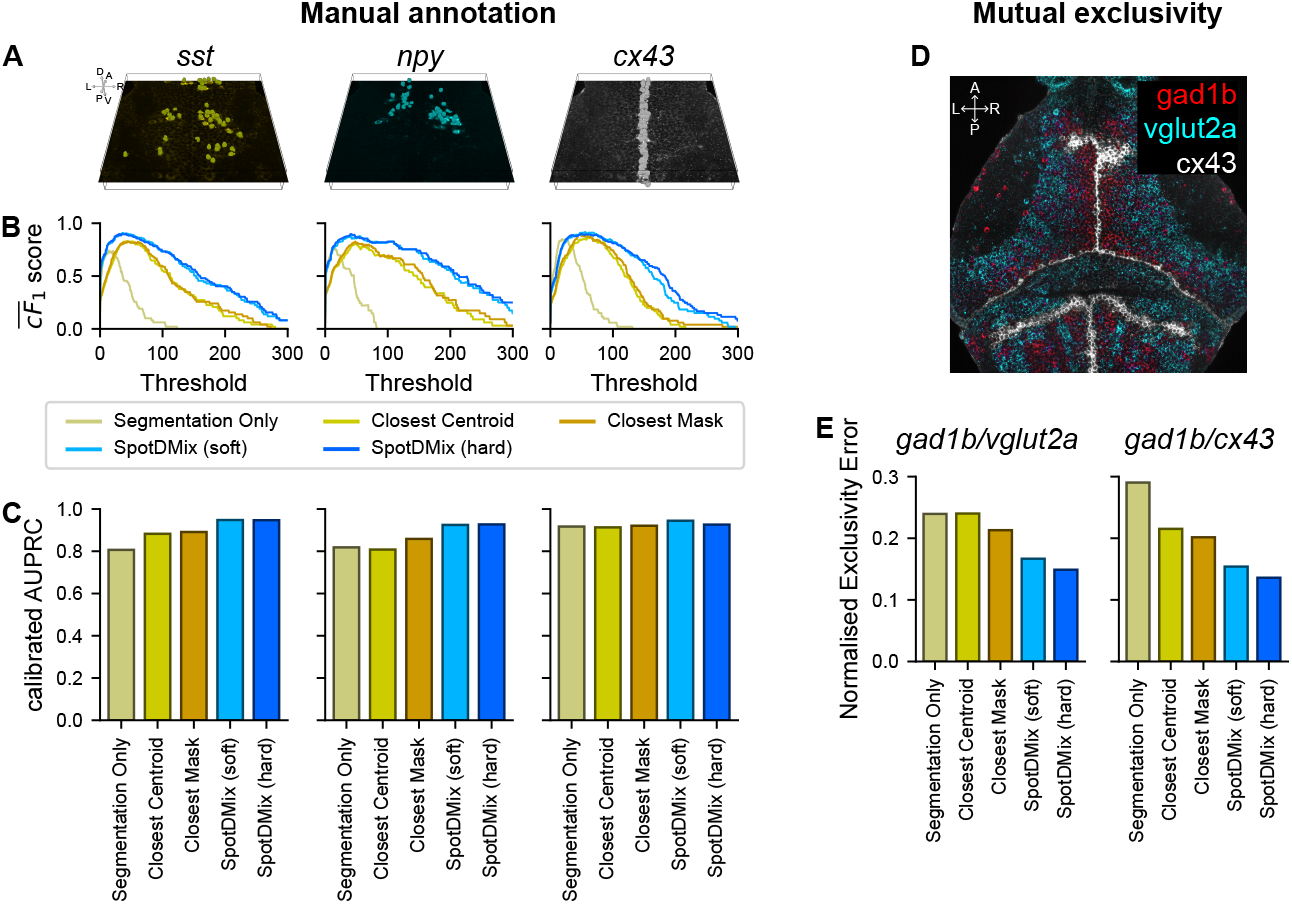
Performance of the different methods on detected spots from larval zebrafish WARP data. **A)** Manual classification of three genes (sst, npy, cx43) in an area of the thalamus (Methods). L/R: Left-/Right. A/P: Anterior/Posterior. D/V: Dorsal/Ventral. Classification performance for the evaluated methods is assessed through the use of **B)** calibrated *F*_1_ curves and **C)** calibrated Area Under the Precision-Recall Curve (AUPRC). **D)** Combined slices of raw fluorescent data from three genes, highlighting mutually exclusive populations of cell types. **E)** The mutual exclusivity in the assigned spot counts is quantified using the normalized Exclusivity Error (Methods) .

Similar to in the first scenario, SpotDMix obtains a significantly higher individual spot assignment accuracy compared to simple methods across the 200 runs (all *p* ≤ 10^−62^, all effect sizes ≥ 1.4, pairwise Wilcoxon rank-sum tests, Suppl. Fig. 3) (Acc._*soft*_ = 72.2 *±* 1.8, Acc._*hard*_ = 77.4 *±* 1.6% versus Acc._SO_ = 70.4 *±* 0.7%, Acc._CC_ = 56.6 *±* 0.9%, Acc._CM_ = 61.9 *±* 0.8%; mean *±* standard deviation) (Fig. 4E). As expected, the accuracy of the mask-based methods is comparable to that of the first scenario as these methods are not influenced by spatial structure. Despite SpotDMix being affected by the mixing of equally likely components as mentioned above, it still outperforms the compared methods on an individual spot level.

The large mixing populations do cause an effect for all methods on the RMSE. This is also where the distinction between the *soft* and *hard* SpotDMix variants becomes clear, as *soft* assigned counts (RMSE_*soft*_ = 3.2 *±* 0.30) are significantly closer to the ground truth data than the *hard* counts are (RMSE_*hard*_ = 4.9 *±* 0.5) (*p* ≤ 10^−66^, effect size = 4.2, Wilcoxon rank-sum test). By not forcing a decision between different cells but instead splitting the count between the cells based on the assignment probabilities *w*_*ik*_, counts are distributed more in line with the ground truth data. Moreover, forcing this decision in the case of *hard* assignments results in a noticeable but moderate level of overfitting for cells with many spots (Fig. 4A).

Both assignment variants on yield significantly more accurate spot counts compared to the simple methods (all *p* ≤ 10^−66^, all effect sizes ≥ 8.6, pairwise Wilcoxon rank-sum tests, Suppl. Fig. 3) (RMSE_SO_ = 20.2 *±* 1.0, RMSE_CC_ = 10.1 *±* 0.5, RMSE_CM_ = 8.6 *±* 0.4) (Fig. 4F).

In an ideal case, having two mutually exclusive populations would mean for a cell with a sufficiently high spot count for gene A to have a spot count of zero for gene B, and vice versa. The deviation from this optimum is measured using a metric termed the (normalized) Exclusivity Error (see Methods). In essence, it is the RMSE in spot counts in cells of the other cell type, normalized by dividing through the standard deviation in spot counts of that cell type.

In this surrogate experiment, both SpotDMix variants obtain a significantly lower exclusivity error than the compared methods (all *p* ≤10^−66^, all effect sizes ≥ 20.0, pairwise Wilcoxon rank-sum tests, Suppl. Fig. 3) (nExcl. Err._*soft*_ = 0.052 *±* 0.006, nExcl. Err._*hard*_ = 0.029 *±* 0.005 versus nExcl. Err._SO_ = 0.252 *±* 0.014, nExcl. Err._CC_ = 0.326 *±* 0.014, nExcl. Err._CM_ = 0.277 *±* 0.012) (Fig. 4D) indicating that it is able to retain a higher exclusivity between the two genes. This is also visually clear in Figure 4C, as the normalized Exclusivity Error directly corresponds to the deviation from either axis. Where before the decisiveness of *hard* assignments from SpotDMix resulted in a larger RMSE, this same decisiveness is what yields it a significantly lower exclusivity error compared to the *soft* assignments (*p* ≤ 10^−65^, effect size = 4.6, Wilcoxon rank-sum test).

This difference is also observed in the classification performance. Here, *cell type* classification is performed instead of the positive/negative classification in the previous scenario (see Methods). Briefly, cells are grouped into four categories for a given threshold: Negative for both, positive for type A, positive for type B, positive for both. Where before low thresholds yielded a high recall, this no longer holds when cells can be classified as being positive for both cell types. The 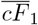 score is now taken as the mean calibrated *F*_1_ score between the classification performance for the two cell types individually (Fig. 4G).

For both variants of SpotDMix the 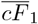 curves almost exclusively exceed the curves of the compared methods. For low and for high thresholds, the *hard* SpotDMix variant achieves a higher 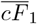 score compared to its *soft* counterpart. However, at medium thresholds the *soft* variant performs slightly better on average. The total classification performance is again summarized using the calibrated AUPRC score, where both SpotDMix variants achieve a significantly higher score across the repeated runs (cAUPRC_*soft*_ = 0.924 *±* 0.007, cAUPRC_*hard*_ = 0.939 *±* 0.006) compared to simple methods (cAUPRC_*SO*_ = 0.817 *±* 0.010, cAUPRC_*CC*_ = 0.777 *±* 0.012, cAUPRC_*CM*_ = 0.802 *±* 0.013) (all p≤ 10^−66^, all effect sizes ≥ 12.5, pairwise Wilcoxon rank-sum tests, Suppl. Fig. 3) (Fig. 4H), agreeing with the 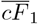 curves and indicating a better classification performance.

### SpotDMix extends well to zebrafish data

To test whether the improved performance of SpotDMix is not only present in surrogate data but also when applied to experimental mRNA transcript data, the classification and exclusivity performance of the different methods is assessed on detected transcripts from larval zebrafish WARP data, using both manually annotated data for three visually clearly expressing genes and whole-brain data for three genes which are characterized as mutually exclusive.

#### Manually annotated data

Positive/negative classification on WARP data is validated through manually annotated data for select genes in a section of the thalamus where many different cell types occur close together ( ∼ 21 *×* 187 *×* 187 *µm*; 2683 cells; see Methods). Cells are classified by the different methods through means of setting a threshold on the assigned counts similar to scenario i). For this, SpotDMix was run on the whole brain data set, and counts are extracted for this smaller section.

For the sparse but highly expressing neuropeptide somatostatin (sst), 95 cells were manually identified as positive (Fig. 5A). Here, both SpotDMix variants obtain a consistently higher *F*_1_ score compared to simple methods for all thresholds (Fig. 5B) and higher cAUPRC scores (cAUPRC_*soft*_ = 0.948, cAUPRC_*hard*_ = 0.947 versus cAUPRC_*SO*_ = 0.807, cAUPRC_*CC*_ = 0.883, cAUPRC_*CM*_ = 0.891) (Fig. 5C), with no clear difference between *soft* and *hard* assignments both in the *F*_1_ curves as in the cAUPRC scores.

Similar behavior is seen for the other neuropeptide-associated and sparsely expressing gene neuropeptide Y (npy) (Fig. 5A), for which 63 cells were labeled as positive. Just as for sst, both SpotDMix variants perform comparable and achieve higher *F*_1_ scores (Fig. 5B) and higher cAUPRC scores compared to simple methods (cAUPRC_*soft*_ = 0.925, cAUPRC_*hard*_ = 0.927 versus cAUPRC_*SO*_ = 0.808, cAUPRC_*CC*_ = 0.859, cAUPRC_*CM*_ = 0.819) (Fig. 5C). Coinciding with the results from surrogate scenario i) with sparse positive cells, using the Closest Mask approach in all cases yields a higher classification performance than Segmentation Only, although the accuracy of the individual assignments is unknown.

The third manually annotated gene is the spatially well-organized and visually distinct gene connexin 43 (cx43), which labels glial cells and is expressed in long but narrow bands along the midline of the fish (Fig. 5A, D). Here, 99 cells in the section were manually identified to be positive (Fig. 5A). Both variants of SpotDMix still obtain higher *F*_1_ scores for nearly all thresholds (Fig. 5B), although the difference at the peak is not as large as for the other evaluated genes. This result is also reflected in the cAUPRC scores, as all obtained values lie closer together with both SpotDMix variants remaining on top (cAUPRC_*soft*_ = 0.945, cAUPRC_*hard*_ = 0.926 versus cAUPRC_*SO*_ = 0.917, cAUPRC_*CC*_ = 0.913, cAUPRC_*CM*_ = 0.921) (Fig. 5C).

#### Genetically defined populations

As discussed in scenario ii), there exist numerous genetically defined populations which are mutually exclusive. Here, the exclusivity of the different assignment methods is assessed between the genes gad1b/vglut2a (inhibitory/excitatory split) and gad1b/cx43 ((inhibitory) neurons/glia split), showcased in Figure 5D. Again, as the different assignment methods can assign a different number of spots each, the normalized Exclusivity Error is used to measure the deviation from mutually exclusive gene counts (Methods).

Similar patterns are observed in both analyzed cases, with *hard* SpotDMix obtaining the lowest error followed closely by its *soft* counterpart, both achieving a better score than simple methods for the gad1b/vglut2a split (nExcl. Err._*soft*_ = 0.167, nExcl. Err._*hard*_ = 0.149 versus nExcl. Err._SO_ = 0.240, nExcl. Err._CC_ = 0.240, nExcl. Err._CM_ = 0.213) as well as for the gad1b/cx43 split (nExcl. Err._*soft*_ = 0.154, nExcl. Err._*hard*_ = 0.136 versus nExcl. Err._SO_ = 0.290, nExcl. Err._CC_ = 0.215, nExcl. Err._CM_ = 0.202) (Fig. 5E).

As in the second surrogate scenario, the decisiveness of *hard* assignments lead to a lower exclusivity error than is obtained with *soft* assignments. Unlike in the surrogate scenario, assigning spots based on the nearest segment with Closest Mask yields a lower exclusivity error than when limiting spot assignment to the segments themselves with Segmentation Only.

## CONCLUSION AND DISCUSSION

Recent advancements in image-based spatial transcriptomics have extended high resolution mRNA transcript imaging to densely packed tissue where the inter-cell distance is much smaller than the width of a single nucleus, making assignment of detected mRNA transcripts to segmented nuclei a complex computational problem. Whereas before simple methods often sufficed, when cell packing is dense Simple methods of assignment often fail to correctly operate in this new regime, leading to patterns where spots are incorrectly assigned to neighboring cells instead. This in turn leads to increased issues with subsequent analysis, as it makes it more challenging to correctly identify cell types at single-cell accuracy and loses information found within the varying expression levels of single genes.

SpotDMix was developed to address these issues by exploiting visually identifiable patterns in the data, with the goal of improving the shortcomings of simple methods and providing a more robust and reliable tool that fits well within a data analysis pipeline. Both the *soft* and *hard* assignment variants of SpotDMix outperform the discussed mask-based methods. This holds not only in the two surrogate scenarios presented here but also extends to the true experimental data set.

The mathematical framework underlying SpotDMix is easily extendable to allow for more complex assignment methods. Several natural extensions are discussed in the section “Future directions” below.

Obtaining accurate genetic profiles of spatially identified cells is essential for studying the complex interplay between different genetically defined cell types, for example when combined with functional recordings (Lovett-Barron et al., 2020; Bugeon et al., 2022). Within the growing field of spatial transcriptomics, SpotDMix leads the way to single-cell accurate assignments of mRNA transcripts in tissue with high cellular density, specifically for cases where the number of genes imaged is limited. SpotDMix is the first spot assignment method to be specifically designed for use in high cellular density tissue. While this work focused on neural data obtained from the larval zebrafish, the underlying principles and methods used are broadly applicable to other tissues with similar characteristics (e.g. in Drosophila).

### Model comparison

Running SpotDMix remains fast and efficient even at whole-brain scale, with the primary bottleneck being the fitting of ellipsoids which itself is highly parallelizable and only needs to be performed once. The relatively simple method of assigning spots to the nearest mask, while also being highly parallelizable, is computationally more intensive to run. A rough overview of the computation times associated with the different methods is provided in Supplementary Table 1.

Deciding on assignments by using the highest probability in the case of the *hard* assignments generally gives a stronger classification boundary between positive and negative cells, leading to a lower mutual exclusivity error in the case of mutually exclusive cells. However, using *soft* assignments generally leads to more accurate spot counts per cell and less overfitting for strongly expressing cells. Both variations have their merits and considerations when it comes to choosing one over the other, specifically in large spatially structured populations of expressing cells. In the case of sparsely expressing cells, such as in surrogate scenario i) or in the neuropeptide-associated genes sst and npy, there is no discernable difference between the two methods.

For surrogate data, the performance of mask-based methods depends highly on the parameters used for generating the data. In the scenarios presented here, Segmentation Only consistently achieved a higher individual spot assignment accuracy than Closest Mask. However, this follows directly from the spot generation parameters, as they can be tweaked just as well as to make Closest Mask more accurate than Segmentation Only by generating more spots outside of the mask. From the performance assessments on zebrafish data, both methods perform comparably with no clear better option. That being said, Closest Mask assigns a significantly larger portion of spots, making the found distributions better match the scale of counts in the experimental data but at a comparable accuracy.

### Data limitations

Informed methods as the one presented here, more than simple mask-based methods, are strongly affected by the quality of the segmentation and spot localization. SpotDMix relies on accurate segmentation to model expected expression patterns around cells, which can yield improper assignments when the fitted ellipsoids do not match the biological counterpart. While any improperly segmented cell is not directly suited for use in further analyzes, SpotDMix can potentially suffer more as spots may still be assigned to this segment instead of to others.

At the same time, the fitting of the ellipsoids can aid in the detection of improper segments present in the data, as is currently already possible using the Jaccard index indicating how well the ellipsoid matches the segment (Suppl. Fig. 1).

On the side of spot localization, noise will remain a persistent factor when dealing with any biological system. As noise in the data or improper spot detection significantly influence the assignment process, regardless of the assignment method employed, assignment methods employing a form of outlier detection form an interesting topic for further examination.

Even in the case of perfect spot detection methods, from the raw fluorescent data it is clear that not all genes express as visually clear patterns around the nucleus. This is especially true for large populations of a single cell type, as mRNA transcripts are not only localized in the cytoplasm directly around the nucleus but can also occur in neural processes such as in axons or dendrites, possibly extending to other regions. For example, this is clear in genes such as gad1b and vglut2a, where positive regions are easy to see but positive cells are impossible to confidently identify at a single-cell level (Fig. 5D), also making manual annotation of these genes near impossible, further highlighting that the presence of this issue in any potential spot assignment algorithm.

### Future directions

The mathematical framework of Expectation Maximization models can be easily extended, such as with the inclusion of outlier detection. Outliers may be modeled as an additional uniform or background component to which spots can be assigned (Melchior and Goulding, 2018; Frühwirth-Schnatter, 2006). As noted above, mRNA transcripts are not always localized close to the cell they belong to as is the case for neural processes, making them impossible to properly assign to any cell. On top of this, generic noise remains a constant factor in any biological data. In these cases, including outlier detection would allow for only counting the spots that follow the expected spot patterns and discarding the others. This would potentially remove the need for setting a relatively high counts threshold to be confident in cell type identification.

Another possibility is to include the measured brightness (fluorescence intensity) of the transcripts as a sort of importance weighting for each of the spots (Gebru et al., 2016). Low intensity spots are more likely to be noise and should contribute less to the weight and shape parameters for a cell. In the current data set, a threshold is set on the intensity to discard low intensity spots, with all spots that are kept being treated as equal in this model. Incorporating a weighted variant of EM could in theory work hand in hand with outlier detection, although it also has the chance to be sensitive to the hyperparameters used.

At the moment, segmented masks are approximated as ellipsoids as this makes the computations fast and as it allows for easily updating the shape parameters of the distribution. However, as the assignment and weight parameters *w*_*ik*_ and *ρ*_*k*_ do not depend on the specific form of the probability distribution, it is possible to use a more complex, static, probability distribution with no iterative shape parameters. For example, it would be possible to directly use the segmented masks to define a unique probability for every voxel near each mask, similar to the way spots are generated in the surrogate scenarios.

We hypothesize, however, that the chosen spatial distributions only need to roughly match the observed distributions in order to achieve accurate assignments, as it is mainly the demixing part of SpotDMix which is responsible for separating positive from negative cells and not the choice of distribution.

When performing outlier detection or when using more complex probability distributions, it could be beneficial to constrain certain parameters through the addition of a prior onto the likelihood function (Bishop, 2006). Even in the current model this can already present useful, as adding a prior on the cell weights *ρ*_*k*_ could limit the number of spots that get assigned to any one cell, potentially providing a solution to the slight overfitting that is seen in the *hard* variant of SpotDMix.

In conclusion, SpotDMix as it stands already leads to improved genetic profiling of cells compared to simple methods, and there is no shortage of ways in which the mathematical framework can be extended towards more complex use cases.

## CODE AVAILABILITY

SpotDMix is available as a Python implementation at https://github.com/Kepser/SpotDMix.

## ACKNOWLEDGMENTS

This work was supported by the NWO VIDI grant 016.Vidi.189.052.

## AUTHOR CONTRIBUTIONS

**K.S**. Conceptualization, Methodology, Software, Formal analysis, Data Curation, Visualization, Writing - Original draft. **L.H**. Conceptualization, Supervision, Writing - Original draft. **E.M**. Conceptualization, Investigation, Data Curation. **G.F**. Conceptualization, Data Curation. **M.E**. Investigation, Data Curation. **P.T**. Resources, Funding acquisition. **M.A**. Supervision, Resources, Funding acquisition, Project administration. **B.E**. Supervision, Writing - Original draft, Funding acquisition, Project administration.

## METHODS

### EM mixture modeling

The locations of spots is modeled as coming from a mixture of spatial probability density functions around each cell, with the goal of assigning each of the *N* spots to one of the *K* cells/components. From this view there are at least two unknowns that need to be estimated: The individual assignments and the relative weighting of each the cells. Both of these are iteratively found by applying the Expectation Maximization algorithm (Dempster et al., 1977; Bishop, 2006), with the “memberships” modeled through an additional hidden variable *z*, as is common in Gaussian Mixture Models (GMM) (Reynolds, 2009). It is also possible to iteratively estimate (some of) the parameters in the chosen probability distributions.

The algorithm shares many similarities with GMMs, with key differences being the fixed component centers and how the distribution shape parameters are shared across all components.

#### Mathematical formulation

Membership is indicated by a latent variable **z** ∈ 1{, 2, …, *K*} ^*N*^, with *z*_*i*_ = *k* indicating that data point *i* belongs to component *k*. Importantly, each data point can only belong to a single component. Given the membership *z*_*i*_ = *k* and distribution parameters *θ* = {*α, β*}, the data points are assumed to be generated from the distribution 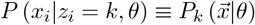.

Bayes’ theorem gives the probability that data point *i* belongs to component *k*:

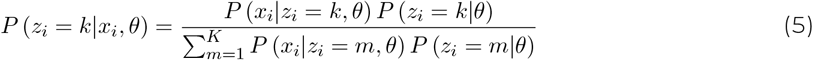

This is termed the ‘*soft* assignment’ and is written as *w*_*ik*_. The prior *P* (*z*_*i*_ = *k* | *θ*) is precisely the relative weight of each component and is denoted as *ρ*_*k*_, obtaining Equation 4. By normalization it holds that 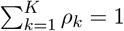 and that 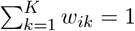

#### EM steps

The log-likelihood of the data is

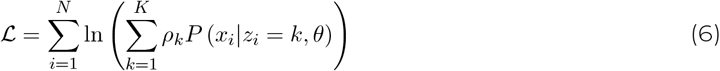

As it is intractable to find the parameters which maximize the complete likelihood function of the data, the EM algorithm maximizes the expectation of the likelihood function given the old parameter estimates. For this log-likelihood the expectation function is

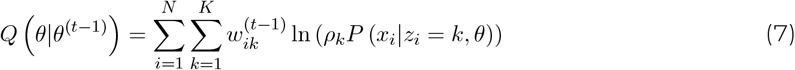

and is maximized by setting the derivatives with respect to *ρ*_*k*_ and *θ* to zero (see below).

Through the updated parameters the new soft assignment probabilities *w*_*ik*_ are calculated and the process repeats until convergence is reached, measured as when the difference between log-likelihoods is below a set threshold: Δℒ *< ε*.

#### Computing radial distributions

A gamma distribution on the transformed distance to the origin *r* (Equation 2) is chosen as the radial density function as spots are expected to occur mostly around the edges of cells with some spread. The exact peak and variance of this distribution are left as shape parameters to be estimated as they can depend on the segmentation and may also differ between genes.

Compared to a regular gamma distribution, the three-dimensional radial gamma distribution was i mple mented with the parameter change *α* → *α* − 2 such that part of the normalization constant 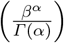 is identical to the one-dimensional case, allowing for easier numerical implementation.

The boundary of each cell *k* is approximated as an ellipsoid specified by a transformation matrix *A*_*k*_ and offset *x*_cell *k*_ such that points on a unit sphere are transformed to the boundary around a cell (see below). For each cell, all nearby spots up to some distance threshold `max_dist` = 10*µm* are transformed using the inverse matrix 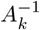 to obtain their transformed distance to the cell center *r* (Eq. 3). In this transformed space a distance of *r* = 1 corresponds to points lying on the boundary of the ellipsoid, with values less than and greater than 1 corresponding to locations inside and outside the ellipsoid respectively. The inclusion of a distance threshold, implemented using sparse matrices, makes running SpotDMix computationally feasible at a brain-wide scale and is a natural addition for spot assignment problems.

The transformed distances are efficiently computed at initialization of SpotDMix and are directly used in the radial gamma distribution. Under the assumption that spots around different cells follow comparable distributions it is now straightforward to define shared shape parameters that are equal for all cells as they lie in the transformed space.

With the exception of an additional normalization term of 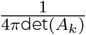 due to the ellipsoidal transformation and the extension of the gamma distribution to three dimensions, it is possible to efficiently implement the probability calculations that are required every iteration using standard libraries such as Python’s scientific computing library `scipy`.

The main processing bottleneck of SpotDMix is the ellipsoid fitting procedure, which is still easily and efficiently be parallelized as every cell segment needs to be processed individually. Further, the obtained preprocessed ellipsoid matrices and offsets can then be used for all gene channels.

For interpretability: The peak of the distribution lies at 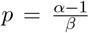 and has a standard deviation of 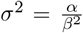. (i.e. spots generated in 3D, distribution of |*r*| has a peak at *p*). For using this function in *d*-dimensional case the substitution *α* → *α* + *d* − 1 should be used for *p* and *σ*^2^.

#### Update functions and parameter initialization

A closed form solution for the mixture weights *ρ*_*k*_ is found through the additional constraint 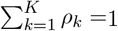 using Lagrange multipliers, yielding 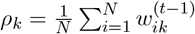 regardless of the specific form of *P*_*k*_ (*x*_*i*_|*θ*). Each *ρ*_*k*_ is exactly updated to be the fraction of spots that are assigned to it’s corresponding component, weighted by the probability of each assignment.

For the use of the radial gamma function the following equations are obtained for updating the shape parameters:

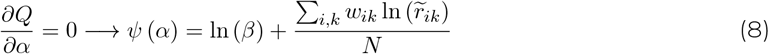

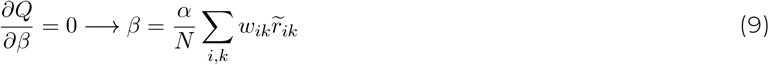

where *ψ* (*x*) is the derivative of the logarithm of the gamma function (so-called digamma function). It is well approximated as *Ψ* (*x*) ≈ ln 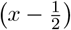 for *x >* 1.5, with the error between the two quickly shrinking for larger *x*. For example, at *x* = 3 there is an error of approximately 0.7%. At *x* = 8 this is reduced to 0.08% and at typical values of *α* of 30 or more, as seen in running SpotDMix in various experiments, the error is further reduced to 0.0014%.

With this approximation in place the exact update functions

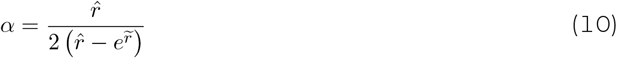

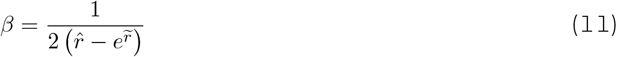

are obtained, where 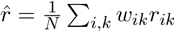 and 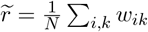 ln (*r*_*ik*_).

To initialize the model *α* and *β* are chosen such that the distribution has a peak at 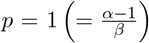 and has a standard deviation of 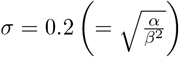, and mixture weights are initialized equally across components: 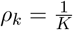 for all *k*.

#### Assignments from probabilities

Turning assignment probabilities into assignments can be done in a variety of ways, the most straightforward of which is what is termed the “*hard* assignment”. Every spot is assigned to the cell to which it has the highest assignment probability. The potential pitfall with this method is that it does not account well for spots that are localized in between two expressing cells. If one of the two cells has a significantly higher mixture weight than the other, all spots in between the two cells are assigned to the one with the higher weight, leading to overfitting of strongly expressing cells and underfitting of lowly expressing cells.

If unique assignments per spot or quantized counts per cell are not a necessity, one can opt for directly using the soft assignment probabilities in determining the spot counts per cell. This way spots are distributed exactly according to their assignment probabilities, circumventing the potential overfitting that may occur with the *hard* assignments. This variant can also be interpreted as the limit of the mean number of spots assigned to each cell when opting for stochastic assignments based on the soft assignment probabilities.

### Ellipsoid approximations

Fitting an ellipsoid to a set of datapoints such that those datapoints form the inside of this ellipsoid is in general a difficult task. Unlike in the 2D case, there exist no closed-form solutions for generalized ellipsoids as they are determined by 9 free parameters (3 for location, 3 for orientation and 3 for size in each direction) that can be represented in a 3×3 symmetric matrix *A* and a size 3 vector ***b*** such that the ellipsoid is described by all points

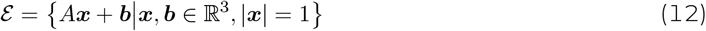

I.e. the transformation from points on a unit sphere using the matrix *A* and offsetting it with ***b***.

There exist numerous iterative methods that efficiently tackle this problem and can find a good fit in just a few iterations (Bertoni, 2010; Allaire et al., 2007; Bektas, 2015). However, for simplicity and efficiency, ellipsoid approximations are done in a single step using the covariance matrix of the locations of the voxels, with the ellipsoid center being the mean of these locations (center of mass). This previously undocumented method works well for voxelized data that is is already roughly ellipsoidal, which generally holds for the segmented cells in the analyzed data set as will be discussed below.

The covariance matrix by itself estimates the shape and orientation of the ellipsoid but with an incorrect size. To find the necessary correcting factor, a theoretical case is examined with negligible grid size with the assumption that the following probability distribution describes the location of the voxels uniformly distributed within the ellipsoid:

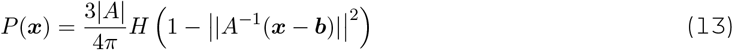

where *H*(***x***) is the Heaviside step function.

Then, using the transformed coordinate space ***x*** = *A****z*** + ***b*** it is follows that

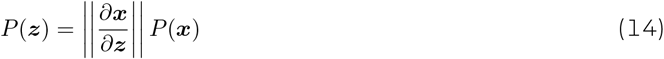

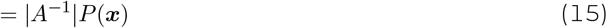

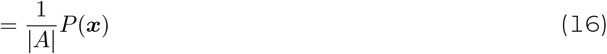

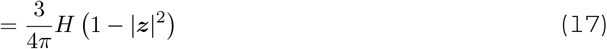

which is now a radially symmetric distribution. Using the spherical coordinates

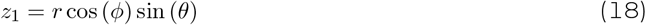

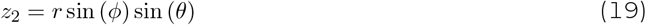

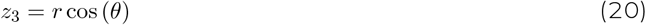

it follows that 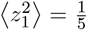. Due to the radial symmetry symmetry, the variance in the three principal directions is identical 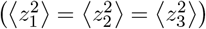, and non-diagonal terms in the covariance matrix are zero. Thus, the covariance matrix corresponding to ***z*** is Cov 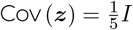 *I* with *I* the identity matrix.

Transforming this back through the use of the linear property of covariance matrices, it follows that

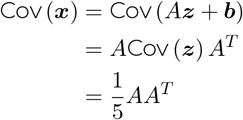

In other words, the covariance matrix corresponding to the voxels ***x*** is a multiple of the product of the transformation matrix *A*. This transformation matrix is then obtained as the Cholevsky decomposition of 5 · Cov (***x***).

To account for irregular spacing of voxels in different axes, ellipsoids are fit in the space corresponding to axis indices, i.e. the space obtained if the spacing would be identical in all directions. The rows of the resulting transformation matrices are then multiplied by the spacing coefficients respectively to obtain a transformation matrix in physical units. This approach slightly underestimates the extent of the segment in the direction with the highest spacing coefficient, although this is negligible for high resolution segments with and extent of tens of voxels in any direction.

The quality of the ellipsoid fit is measured using the Jaccard index. It is a measure of the overlap between the original segment and all the voxels that lie inside of the fitted ellipsoid. A score of 1 indicates a perfect overlap, which in practice is never achieved in part due to the fixed spacing between points. Most of the segments in the volume are well approximated by ellipsoid, as approximately 74.6% of segments has a Jaccard index score greater than 0.8, and 94.3% has a score greater than 0.7 (Suppl. Fig. 1B).

### Performance metrics

#### Accuracy

At an individual spot level, the accuracy is defined as the percentage of spots which is correctly assigned. However, for the *soft* variant of SpotDMix there are no direct assignments, leading to no direct measure of accuracy here.

Despite this, in order to make comparisons with the *soft* variant a different measure of accuracy is defined. As every spot still has a ground truth label associated with it, and as there exist probabilities of assignment for between every spot-cell pair, the accuracy for the *soft* variant of SpotDMix is defined as the overlap of the soft assignment probabilities with the ground truth labels, by taking the mean probability that a spot is assigned to its corresponding ground truth cell:

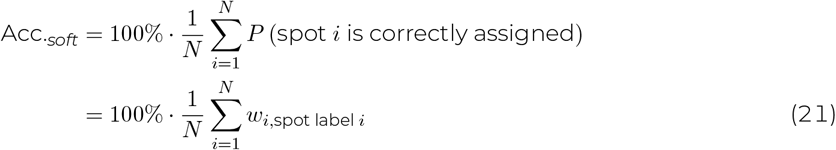

This metric has a similar interpretability as the regular measure of accuracy, and makes it possible to assess the performance of the *soft* variant of SpotDMix at the single-spot level. Naturally, but not exclusively, this accuracy falls below that of the *hard* counterpart as probabilities of assignment can not exceed 1, and probabilities will always be slightly below 1 when there are other cells nearby.

#### Precision and recall

High class imbalance causes high classification accuracy simply due to the overwhelming number of negative samples (Flach and Kull, 2015; Saito and Rehmsmeier, 2015; Siblini et al., 2020). To deal with this, precision and recall are used instead, as they do not depend on the number of True Negatives after classification. More specifically, to make comparisons between different scenarios possible and to combine statistics despite random variations in the number of positive cells, the typical measure of precision is replaced with calibrated precision (Siblini et al., 2020), using a reference value of *π*_0_ = 0.03 for all experiments, corresponding to a scenario where 3% of cells is positive.

Typically, for any threshold, it is a trade-off between the precision and recall, with an ideal classifier achieving a high score for both simultaneously. The *F*_1_ score measures this trade-off by taking the harmonic mean of the two values, 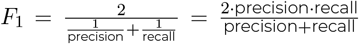. The calibrated *F*_1_ score is obtained by replacing the precision in this function with the calibrated precision, and is used throughout.

For surrogate scenario ii), classification of cell types based on assigned spot counts was done into four categories; Negative for both, positive for type A, positive for type B and positive for both. As there is no distinction in the way the surrogate data is generated between type A and B, a single threshold is used for both spot counts simultaneously. Here, the precision and recall are defined in a one-v-all matter, meaning that positive cases per cell type are counted as usual when the determined classification agrees with the ground truth label for this cell type, and negative cases are counted when the determined classification is any other category. The (calibrated) *F*_1_ scores are calculated for each threshold for both cell types, and are averaged over the two cell types, obtaining the mean calibrated score denoted as 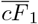.

As the threshold is a free parameter in (*c*)*F*_1_ curves, and as different methods can have differently shaped (*c*)*F*_1_ curves as seen in the surrogate scenarios, it is often difficult to make one-on-one comparisons. To assign a single scalar value to each method, another metric is used that measures the Area Under the Curve (AUC) in the (calibrated) Precision-Recall space (cAUPRC), where individual points correspond to a single threshold and the area summarizes the performance for all thresholds. This area under the curve is measured not with linear interpolation but with rectangles, also sometimes referred to as the Average Precision score, as is more applicable in this non-linear space (Davis and Goadrich, 2006; Flach and Kull, 2015).

#### (Normalized) Exclusivity Error

For data that is expected to be mutually exclusive, the exclusivity error is measured by classifying cells as positive for type A (*C*_*A*_ *> C*_*B*_) or type B (*C*_*B*_ *> C*_*A*_) and exclude cells when |*C*_*A*_ − *C*_*B*_ | *<* 1 (i.e. when cells get assigned a (near) equal amount of spots for both genes), where *C*. is the number of assigned spots (“count”) to a cell for a certain type. Next, for type A cells, the RMSE for type B counts is measured (with expected value 0), and vice versa for type B cells. The exclusivity error is then the weighted mean between these two values, weighted by the total number of cell types per category, and is indicative of the variance in counts for the other cell type.

Different methods can assign a different number of cells. Especially the Segmentation Only variant only assigns a small number of spots, which would naturally yield a lower exclusivity error. To account for this, the normalized Exclusivity Error is introduced by dividing the error by the standard deviation in spot counts for that gene:

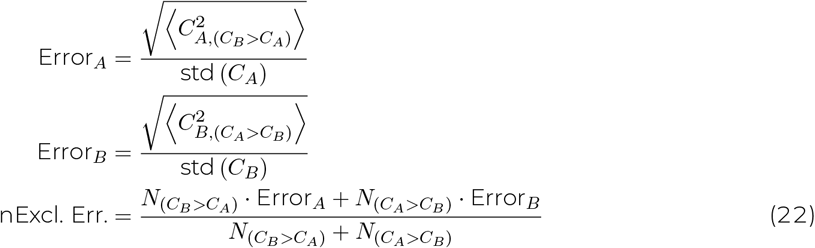

Where ⟨.⟩ denotes the mean of that value, *N*_(.)_ denotes the number of cells in that category, and cells with |*C*_*A*_ − *C*_*B*_| *<* 1 are excluded as mentioned.

### Surrogate data generation

Surrogate spot data was generated in order to more extensively assess and compare the performance of different methods, and also due to the low amount of validated or ground truth data.

Spots are generated on a cell by cell basis matching the segmented cell mask through a Gaussian convolution of the dilated cell boundary, with the number of spots to generate per cell sampled from a gamma distribution similar to the manually annotated data (Suppl. Fig. 2). More precisely, first, the segmented binary mask of a cell is dilated in all directions by `peak_offset` = 4 voxels in the x- and y-direction, and by 2 pixels in the z-direction, based on the scaling factor in this direction. The boundary of this image is taken and convolved with a Gaussian with standard deviation of *σ* = 3.5 in the x- and y-direction, with the standard deviation used in the z-direction again scaled according to the scaling factor corresponding to this direction.

This processed image is then normalized and spots are sampled from voxel locations. The offset of the boundary ensures that the majority of the generated spots lie outside of the cell they belong to, as is also observed in the experimental data, and the Gaussian convolution ensures variability in the distance to the mask of origin.

The parameters used for generating the surrogate data are carefully chosen to reflect the patterns observed in the experimental data as best as possible. It is easy to see that the choice of parameters has a large impact on specifically the performance of the simple methods. Namely, the amount of spots that is generated inside the mask itself and what fraction of spots is generated inside of other cells (by chance), both directly determine an upper limit on the accuracy of the mask-based methods Segmentation Only and Closest Mask. To minimize user bias and to model spot generation as accurately as possible, manually annotated data for the three genes was used to compute several statistics relating to the distribution of spots, including the distance to the nearest mask, the percentage of spots that lies inside of any segments and the percentage of spots that lies inside of positively classified segments.

From an experiment where spots were generated right outside of cell segments (`peak_offset` = 1, *σ* = 0.1) it was found that roughly 40% of space directly neighboring any cell is taken up by other cells. However, in the experimental data set only ∼ 10% of spots is found inside of negatively classified cells. Visual inspection confirmed that the large majority of spots are localized in the space in between cells, likely following biological structures. To account for this, and to not give a strong disadvantage to simple methods, every spot that was generated inside of a neighboring cell was removed based on a carefully calculated probability of 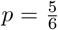. By removing every 5 out of 6 spots from neighboring cells in the generated data, the percentage of spots outside of the original cell but inside of neighboring cells indeed dropped to around 10%.

For each positive cell the number of spots to generate for it (excluding neighbor removal) was sampled from a gamma distribution with a peak at *p* = 70 spots and a standard deviation of *σ* = 40 (Suppl. Fig. 2). Cells that had less than 20 spots after the neighbor overlap removal step were stripped of all their spots, ensuring a clear gap between positively and negatively expressing cells and making sure that all positive cells are also visually identifiable as such.

#### Sparse positive cells

To thoroughly assess the performance of different methods on data similar to that of neuropeptide-associated genes, data sets were generated with cell segmentation taken from on a small 3D volume in the hindbrain ( ∼ 85 *×* 93 *×* 93 *µm*; 5040 cells). Surrogate spots are generated around randomly selected segments.

In the manually annotated data only roughly 3% of sst and npy cells were positively identified on average. In this surrogate scenario, cell labels are randomly assigned with each cell having exactly this probability of being positive. Not only does this model the spatial distribution of neuropeptide-associated genes well, it also presents an idealistic and minimalistic testing ground for spot assignment.

Spots are generated for the cells labeled as positive according to the procedure described above. The performance is assessed in 1000 of these runs to account for variability and the low number of positive cells.

#### Mutually exclusive neighboring bands

To assess the performance of the different methods both for large and neighboring populations, the same area in the hindbrain as above was used for the generation of spots corresponding to two closely bordering populations.

Three-dimensional Perlin noise patterns are used to generate labels for two cell types based on a threshold, as Perlin noise smoothly blends between gentle peaks with a value of 1 and troughs with a value of -1 (Suppl. Fig. 2C). A noise pattern is generated with the same dimensions as the segmented area, and the full band of all positive cells is achieved by looking at the values of the noise pattern at the centroid of each cell. Cells with a value of |noise (*x*_cell_) | *<* 0.1 are selected to be positive, yielding a narrow band that smoothly meanders throughout the volume. Type A/Type B labels are assigned based on the sign of the noise pattern at their locations, giving a fair split between the two cell types due to the symmetry in the values of the Perlin noise.

Again, spots are generated according to the procedure described above for cells labeled as positive for either cell type. After generation, spots are split into two groups based on the type of cell they belong to, as the different groups are processed by the assignment methods individually. Performance is assessed in 200 repeated runs of this scenario.

### Manual annotation

A small area in the thalamus was selected for manual annotation of cells (∼ 21 *×* 187 *×* 187 *µm*; 2683 cells), as this is where several neuropeptide-associated genes such as sst, npy and penkA express close together. The selected area also includes a band of cx43 expressing cells down the midline of the fish corresponding to glial cells. A manual positive/negative labeling was performed for three genes (sst, npy, cx43) as individual spot assignment is often impossible to perform with certainty when there are neighboring positive cells and is in general challenging for many spots. On top of this, the classification performance of different methods is in some aspects a more important metric than individual spot assignment accuracy, though they are highly related and accurate spot counts can reveal intricacies within the data.

The segmentation of all cells contained in this area were loaded into Blender for easy navigation in three dimensions. A horizontal plane was added displaying the raw fluorescent imaging data, which updates to automatically display the fluorescent data for that slice when the plane slides up and down the volume (Fig. 5A). This approach was chosen in favor of displaying the detected spot positions as spot detection and filtering algorithms can sometimes fail or introduce bias towards strongly expressing regions.

Besides the classification into positive and negative groups, cells with clear segmentation issues were also identified and left out in the analyzes. This is done as these masks often do not correspond one-to-one with a cell, making proper classification impossible. The most common of such segments include single cells that are split in half, two cells which are joined together and miscellaneous shapes (Suppl. Fig. 1A). In total 47 segments were identified as having improper segmentation and were left out. This however only includes segmentation issues that were spotted near or at positively expressing regions, as the segmentation issues at further away locations do not influence the classification performance.

However, as segmentation issues remain a part of the large data set, all methods were run without the removal of any segments. It is only in measuring the classification performance for this manually annotated data set that the obtained counts for the identified improper segments were omitted.

### Mask-based assignment methods

Prior FISH methods typically only assigned spots that already overlapped with segmentation (Wang et al., 2021; Chen et al., 2015b) as a large part of the spots occurred there. However, this no longer holds for zebrafish data, where typically only 20 − 40% (depending on the gene) of spots lie within the segmentation. This priorly used method is termed *Segmentation Only* and is also included in this analysis. However, as this method only assigns a small fraction of the total number of spots, the accuracy is calculated only based on the spots it did assign.

Two additional simple, mask-based, methods are evaluated that assign all spots (up to some distance threshold). The first of these is termed the *Closest Centroid* approach. As the center of mass (centroid) of each segmented mask is easily calculated during the segmentation process, the easiest to implement assignment method is assigning spots to the closest cell centroid. While this method does not deal well with varying cell sizes, it can be implemented extremely efficiently using a *k*-d tree (e.g. using `scipy`).

The second additional approach is termed *Closest Mask*. Here, the spots are assigned to the closest segment boundary, with the underlying idea being that the intercellular space can be evenly divided between cells as a proxy for membrane borders. Spot locations are turned into indices for the full segmented mask containing all cells as corresponding indices. Around each spot, a box of predetermined size is extracted and distances are computed for all voxels which correspond to cells in this box. The spot is then assigned to the cell that has the minimum distance from a voxel to the spot. While there are most likely ways to optimize the specific implementation, whilst still being able to run on the whole-brain level, every implementation of this method will remain computationally costly.

### Data acquisition

The larval zebrafish WARP data used in this study was obtained by the Ahrens lab at the Janelia Research Campus (Marquez-Legorreta et al., in prep.). Briefly, a larval zebrafish is expanded using Expansion Microscopy (Chen et al., 2015a; Tillberg et al., 2016) to obtain higher resolution images of the diffraction limited fluorescence at mRNA transcripts, akin to EASI-FISH (Wang et al., 2021). Imaging was done at whole-brain scale for several rounds with up to three genes imaged per round. Cell segments were obtained through a nuclear DAPI stain, labeling cell nuclei, yielding on the order of ∼ 2 · 10^6^ cells in total. The number of spots detected per gene varies largely depending on the gene in question, but can easily exceed ∼ 10^9^ for widely expressing genes such as gad1b or vglut2a. Combined this leads to a large-scale 3-dimensional data set of dense cellular tissue with mRNA transcripts in between, making efficient and automated tools essential for the processing that is required here.

## SUPPLEMENTARY MATERIALS

**Supplementary Figure 1:**
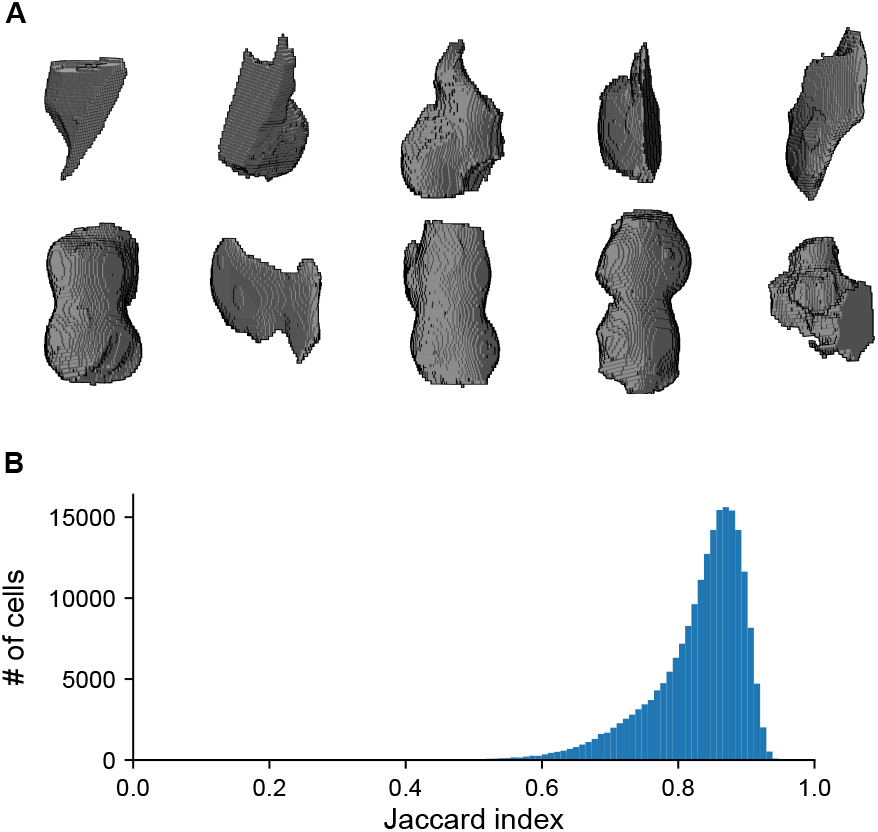
**A)** 10 example segments that were excluded from the manual annotation data, including half cells, double cells and generally odd shapes. **B)** Distribution of Jaccard index scores for all segments in the zebrafish brain indicating the quality of the ellipsoid fit. The Jaccard index applied in this context measures the overlap between the original segment and all voxels that lie inside of the fitted ellipsoid.

**Supplementary Figure 2:**
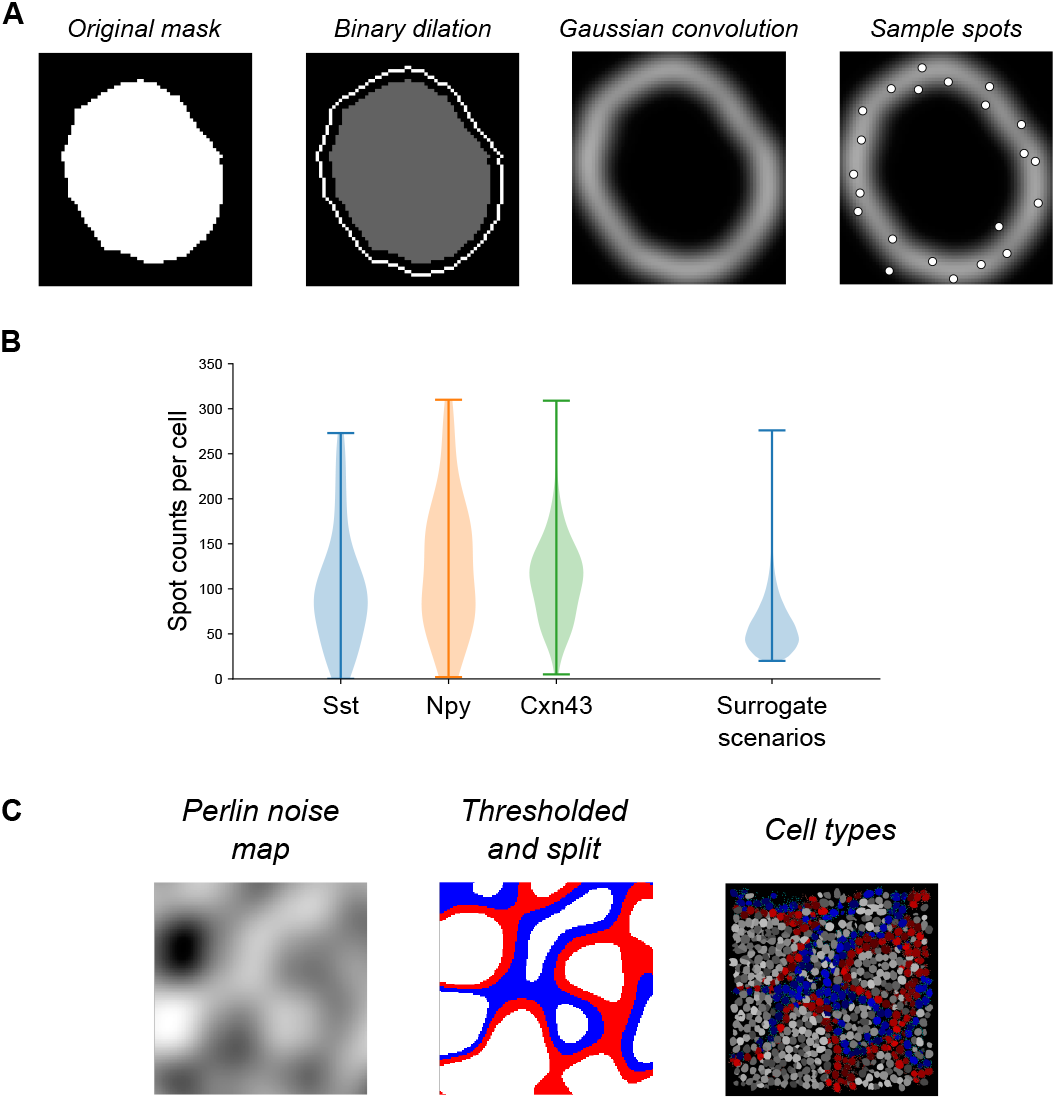
**A)** Steps involved in the generation of spots around segments. Masks are dilated by peak_offset voxels and the boundary of this image is taken. A Gaussian convolution filter is applied to the boundary of the dilated image with standard deviation *σ*. The image is normalized and locations are sampled from this image with replacement. Additional random sub-pixel offsets are applied to remove the perfect grid spacing and cause unique spots when a specific voxel is chosen more than once. **B)** Comparison between manual annotation counts and generated data. Manual annotation counts are taken as the Closest Mask counts corresponding to the cells manually labeled as positive for the three genes shown here. **C)** Process of assigning cell types in surrogate scenario ii). A Perlin noise map is generated and thresholded. Cells are split between two types based on the sign of the noise map at their location.

**Supplementary Figure 3:**
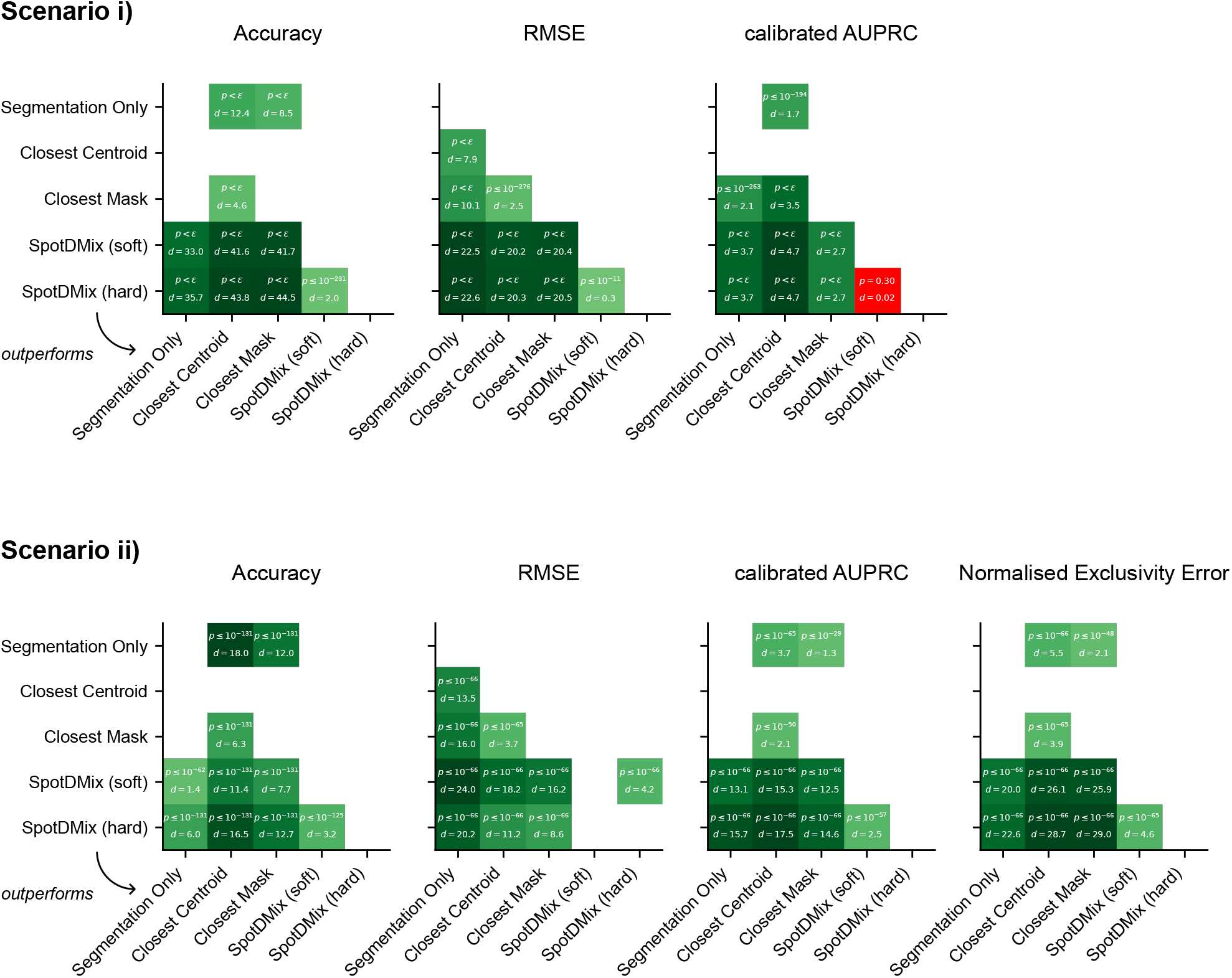
Pairwise *p*-values (one-sided Wilcoxon rank-sum tests) and effect sizes (Cohen’s *d*) between the compared methods for the used metrics in the two surrogate scenarios. Each shown square represents the method on the left outperforming (on average) the method on the bottom. Not significant *p*-values are indicated in red squares. *ε* indicates a *p*-value below the 64-bit floating point precision used in the calculation, which can conservatively be estimated to be less than 10^−307^.

**Supplementary Table 1:**
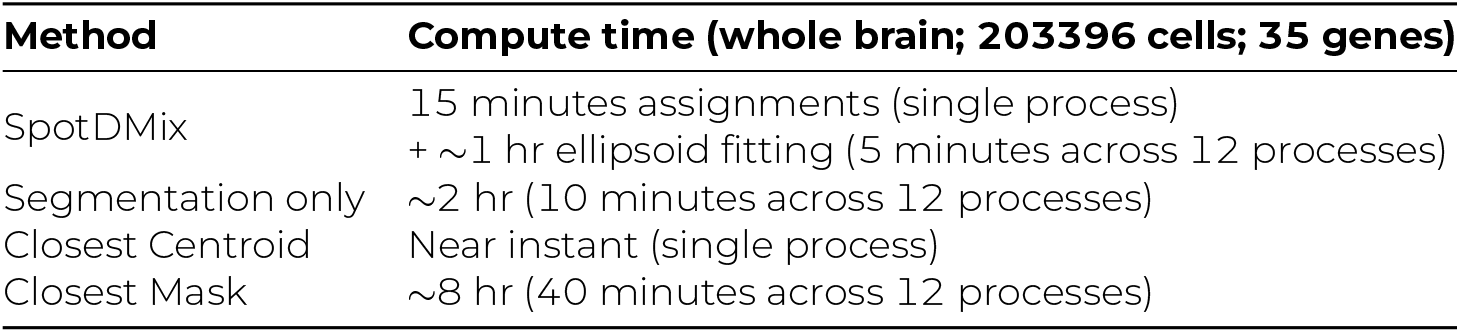
Rough comparison of the computational costs of the different methods compared in this study when ran on the whole-brain larval zebrafish EASI-FISH data set. Parallel processing was done on the Compute Cluster at Janelia Research Campus. Both *soft* and *hard* gene counts are calculated simultaneously at no additional cost.

## REFERENCES

M. B. Ahrens, M. B. Orger, D. N. Robson, J. M. Li, and P. J. Keller. Whole-brain functional imaging at cellular resolution using light-sheet microscopy. Nature Methods, 10(5):413–420, 05 2013. ISSN 1548-7091, 1548-7105. URL 10.1038/nmeth.2434.

S. Allaire, J.-J. Jacq, V. Burdin, and C. Roux. Ellipsoid-constrained robust fitting of quadrics with application to the 3d morphological characterization of articular surfaces. In 2007 29th Annual International Conference of the IEEE Engineering in Medicine and Biology Society, pages 5087–5090. IEEE, 2007. URL 10.1109/IEMBS.2007.4353484.

A. Andersson, A. Behanova, C. Avenel, J. Windhager, F. Malmberg, and C. Wählby. Points2regions: Fast, interactive clustering of imaging-based spatial transcriptomics data. bioRxiv, 2024. URL 10.1101/2022.12.07.519086.

L. Appelbaum, G. X. Wang, G. S. Maro, R. Mori, A. Tovin, W. Marin, T. Yokogawa, K. Kawakami, S. J. Smith, Y. Gothilf, et al. Sleep–wake regulation and hypocretin–melatonin interaction in zebrafish. Proceedings of the National Academy of Sciences, 106(51):21942–21947, 2009. URL 10.1073/pnas.906637106.

S. Bektas. Least squares fitting of ellipsoid using orthogonal distances. Boletim de ciências geodésicas, 21:329–339, 2015. URL 10.1590/S1982-21702015000200019.

B. Bertoni. Multi-dimensional ellipsoidal fitting. Department of Physics, South Methodist University, Tech. Rep. SMU-HEP-10-14, 2010.

C. M. Bishop. Pattern recognition and machine learning, chapter Mixture Models and EM, page 423–455. Springer New York, 2006.

S. Bugeon, J. Duffield, M. Dipoppa, A. Ritoux, I. Prankerd, D. Nicoloutsopoulos, D. Orme, M. Shinn, H. Peng, H. Forrest, A. Viduolyte, C. B. Reddy, Y. Isogai, M. Carandini, and K. D. Harris. A transcriptomic axis predicts state modulation of cortical interneurons. Nature, 607(7918):330–338, 07 2022. ISSN 0028-0836, 1476-4687. URL 10.1038/s41586-022-04915-7.

F. Chen, P. W. Tillberg, and E. S. Boyden. Expansion microscopy. Science, 347(6221):543–548, 2015a. URL 10.1126/science.1260088.

K. H. Chen, A. N. Boettiger, J. R. Moffitt, S. Wang, and X. Zhuang. Spatially resolved, highly multiplexed rna profiling in single cells. Science, 348(6233):aaa6090, 2015b. URL 10.1126/science.aaa6090.

S. Codeluppi, L. E. Borm, A. Zeisel, G. La Manno, J. A. Van Lunteren, C. I. Svensson, and S. Linnarsson. Spatial organization of the somatosensory cortex revealed by osmFISH. Nature Methods, 15(11):932–935, 11 2018. ISSN 1548-7091, 1548-7105. URL 10.1038/s41592-018-0175-z.

N. Crosetto, M. Bienko, and A. van Oudenaarden. Spatially resolved transcriptomics and beyond. Nature Reviews Genetics, 16(1):57–66, 01 2015. ISSN 1471-0064. URL 10.1038/nrg3832.

J. Davis and M. Goadrich. The relationship between precision-recall and roc curves. In Proceedings of the 23rd international conference on Machine learning, pages 233–240, 2006. URL 10.1145/1143844.1143874.

A. P. Dempster, N. M. Laird, and D. B. Rubin. Maximum Likelihood from Incomplete Data Via the EM Algorithm. Journal of the Royal Statistical Society: Series B (Methodological), 39(1):1–22, 09 1977. ISSN 00359246. URL 10.1111/j.2517-6161.1977.tb01600.x.

P. Flach and M. Kull. Precision-recall-gain curves: Pr analysis done right. Advances in neural information processing systems, 28, 2015.

S. Frühwirth-Schnatter. Finite Mixture and Markov Switching Models. Springer Series in Statistics. Springer New York, 2006. ISBN 978-0-387-32909-3. URL 10.1007/978-0-387-35768-3.

I. D. Gebru, X. Alameda-Pineda, F. Forbes, and R. Horaud. EM Algorithms for Weighted-Data Clustering with Application to Audio-Visual Scene Analysis. IEEE Transactions on Pattern Analysis and Machine Intelligence, 38(12):2402–2415, 12 2016. ISSN 0162-8828, 2160-9292. URL 10.1109/TPAMI.2016.2522425.

H. Herzog. 30 years of npy research. Neuropeptides, 46(6):251, 2012. ISSN 0143-4179. URL 10.1016/j.npep.2012.10.002.30 years of NPY research.

M. Lovett-Barron, A. S. Andalman, W. E. Allen, S. Vesuna, I. Kauvar, V. M. Burns, and K. Deisseroth. Ancestral Circuits for the Coordinated Modulation of Brain State. Cell, 171(6):1411–1423.e17, 11 2017. ISSN 00928674. URL 10.1016/j.cell.2017.10.021.

M. Lovett-Barron, R. Chen, S. Bradbury, A. S. Andalman, M. Wagle, S. Guo, and K. Deisseroth. Multiple convergent hypothalamus–brainstem circuits drive defensive behavior. Nature Neuroscience, 23(8):959–967, 08 2020. ISSN 1097-6256, 1546-1726. URL 10.1038/s41593-020-0655-1.

S. Mages, N. Moriel, I. Avraham-Davidi, E. Murray, J. Watter, F. Chen, O. Rozenblatt-Rosen, J. Klughammer, A. Regev, and M. Nitzan. TACCO unifies annotation transfer and decomposition of cell identities for single-cell and spatial omics. Nature Biotechnology, 02 2023. ISSN 1087-0156, 1546-1696. URL 10.1038/s41587-023-01657-3.

P. Mahou, J. Vermot, E. Beaurepaire, and W. Supatto. Multicolor two-photon light-sheet microscopy. Nature Methods, 11(6):600–601, 06 2014. ISSN 1548-7105. URL 10.1038/nmeth.2963.

J. C. Marques, M. Li, D. Schaak, D. N. Robson, and J. M. Li. Internal state dynamics shape brain-wide activity and foraging behaviour. Nature, 577(7789):239–243, 01 2020. ISSN 0028-0836, 1476-4687. URL 10.1038/s41586-019-1858-z.

E. Marquez-Legorreta, G. M. Fleishman, L. W. Hesselink, M. Eddison, K. Smeets, C. Stringer, P. Keller, S. Narayan, A. B. Chen, S. M. Sternson, B. Englitz, P. W. Tillberg, and M. B. Ahrens. Whole-brain co-mapping of gene expression and neuronal activity at cellular resolution in behaving zebrafish. in prep.

P. Melchior and A. Goulding. Filling the gaps: Gaussian mixture models from noisy, truncated or incomplete samples. Astronomy and Computing, 25:183–194, 10 2018. ISSN 22131337. URL 10.1016/j.ascom.2018.09.013.

J. R. Moffitt, E. Lundberg, and H. Heyn. The emerging landscape of spatial profiling technologies. Nature Reviews Genetics, 23(12):741–759, 12 2022. ISSN 1471-0056, 1471-0064. URL 10.1038/s41576-022-00515-3.

J. Park, W. Choi, S. Tiesmeyer, B. Long, L. E. Borm, E. Garren, T. N. Nguyen, B. Tasic, S. Codeluppi, T. Graf, M. Schlesner, O. Stegle, R. Eils, and N. Ishaque. Cell segmentation-free inference of cell types from in situ transcriptomics data. Nature Communications, 12(1):3545, 06 2021. ISSN 2041-1723. URL 10.1038/s41467-021-23807-4.

G. Partel and C. Wählby. Spage2vec: Unsupervised representation of localized spatial gene expression signatures. The FEBS Journal, 288(6):1859–1870, 03 2021. ISSN 1742-464X, 1742-4658. URL 10.1111/febs.15572.

V. Petukhov, R. J. Xu, R. A. Soldatov, P. Cadinu, K. Khodosevich, J. R. Moffitt, and P. V. Kharchenko. Cell segmentation in imaging-based spatial transcriptomics. Nature Biotechnology, 40 (3):345–354, 03 2022. ISSN 1087-0156, 1546-1696. URL 10.1038/s41587-021-01044-w.

S. Prabhakaran. Sparcle: Assigning transcripts to cells in multiplexed images. Bioinformatics Advances, 2(1):vbac048, 01 2022. ISSN 2635-0041. URL 10.1093/bioadv/vbac048.

D. Reynolds. Gaussian Mixture Models. In S. Z. Li and A. Jain, editors, Encyclopedia of Biometrics, pages 659–663. Springer US, 2009. ISBN 978-0-387-73002-8. URL 10.1007/978-0-387-73003-5_196.

T. Saito and M. Rehmsmeier. The precision-recall plot is more informative than the roc plot when evaluating binary classifiers on imbalanced datasets. PloS one, 10(3):e0118432, 2015. URL 10.1371/journal.pone.0118432.

A. K. Sandoval-Talamantes, B. Gómez-González, D. Uriarte-Mayorga, M. Martínez-Guzman, K. A. Wheber-Hidalgo, and A. Alvarado-Navarro. Neurotransmitters, neuropeptides and their receptors interact with immune response in healthy and psoriatic skin. Neuropeptides, 79:102004, 2020. URL 10.1016/j.npep.2019.102004.

Siblini, J. Fréry, L. He-Guelton, F. Oblé, and Y.-Q. Wang. Master your metrics with calibration. In M. R. Berthold, A. Feelders, and G. Krempl, editors, Advances in Intelligent Data Analysis XVIII, pages 457–469, Cham, 2020. Springer International Publishing. URL 10.1007/978-3-030-44584-3_36.

L. Swanson and P. E. Sawchenko. Hypothalamic integration: organization of the paraventricular and supraoptic nuclei. Annual review of neuroscience, 6(1):269–324, 1983. URL 10.1146/annurev.ne.06.030183.001413.

P. W. Tillberg, F. Chen, K. D. Piatkevich, Y. Zhao, C.-C. Yu, B. P. English, L. Gao, A. Martorell, H.-J. Suk, F. Yoshida, et al. Protein-retention expansion microscopy of cells and tissues labeled using standard fluorescent proteins and antibodies. Nature biotechnology, 34(9):987–992, 2016. URL 10.1038/nbt.3625.

X. Wang, W. E. Allen, M. A. Wright, E. L. Sylwestrak, N. Samusik, S. Vesuna, K. Evans, C. Liu, C. Ramakrishnan, J. Liu, G. P. Nolan, F.-A. Bava, and K. Deisseroth. Three-dimensional intact-tissue sequencing of single-cell transcriptional states. Science, 361(6400):eaat5691, 07 2018. ISSN 0036-8075, 1095-9203. URL 10.1126/science.aat5691.

Y. Wang, M. Eddison, G. Fleishman, M. Weigert, S. Xu, T. Wang, K. Rokicki, C. Goina, F. E. Henry, A. L. Lemire, U. Schmidt, H. Yang, K. Svoboda, E. W. Myers, S. Saalfeld, W. Korff, S. M. Sternson, and P. W. Tillberg. Easi-fish for thick tissue defines lateral hypothalamus spatio-molecular organization. Cell, 184(26):6361–6377.e24, 2021. ISSN 0092-8674. URL 10.1016/j.cell.2021.11.024.

S. Xu, H. Yang, V. Menon, A. L. Lemire, L. Wang, F. E. Henry, S. C. Turaga, and S. M. Sternson. Behavioral state coding by molecularly defined paraventricular hypothalamic cell type ensembles. Science, 370(6514):eabb2494, 10 2020. ISSN 0036-8075, 1095-9203. URL 10.1126/science.abb2494.

C. Yanicostas, E. Barbieri, M. Hibi, A. Brice, G. Stevanin, and N. Soussi-Yanicostas. Requirement for zebrafish ataxin-7 in differentiation of photoreceptors and cerebellar neurons. PloS one, 7(11): e50705, 2012. URL 10.1371/journal.pone.0050705.

